# Biogenesis of P-TEFb in CD4^+^ T cells to reverse HIV latency is mediated by protein kinase C (PKC)-independent signaling pathways

**DOI:** 10.1101/2021.04.26.441433

**Authors:** Uri Mbonye, Konstantin Leskov, Meenakshi Shukla, Saba Valadkhan, Jonathan Karn

**Author notes:** Corresponding Authors (UM); (JK).

## Abstract

The switch between HIV latency and productive transcription is regulated by an auto-feedback mechanism initiated by the viral trans-activator Tat, which functions to recruit the host transcription elongation factor P-TEFb to proviral HIV. A heterodimeric complex of CDK9 and one of three cyclin T subunits, P-TEFb is expressed at vanishingly low levels in resting memory CD4^+^ T cells and cellular mechanisms controlling its availability are central to regulation of the emergence of HIV from latency. Using a well-characterized primary T-cell model of HIV latency alongside healthy donor memory CD4^+^ T cells, we characterized specific T-cell receptor (TCR) signaling pathways that regulate the generation of transcriptionally active P-TEFb, defined as the coordinate expression of cyclin T1 and phospho-Ser175 CDK9. Protein kinase C (PKC) agonists, such as ingenol and prostratin, stimulated active P-TEFb expression and reactivated latent HIV with minimal cytotoxicity, even in the absence of intracellular calcium mobilization with an ionophore. Unexpectedly, inhibition-based experiments demonstrated that PKC agonists and TCR-mobilized diacylglycerol signal through MAP kinases ERK1/2 rather than through PKC to effect the reactivation of both P-TEFb and latent HIV. Single-cell and bulk RNA-seq analyses revealed that of the four known isoforms of the Ras guanine nucleotide exchange factor RasGRP, RasGRP1 is by far the predominantly expressed diacylglycerol-dependent isoform in CD4^+^ T cells. RasGRP1 should therefore mediate the activation of ERK1/2 via Ras-Raf signaling upon TCR co-stimulation or PKC agonist challenge. Combined inhibition of the PI3K-mTORC2-AKT-mTORC1 pathway and the ERK1/2 activator MEK prior to TCR co-stimulation abrogated active P-TEFb expression and substantially suppressed latent HIV reactivation. Therefore, contrary to prevailing models, the coordinate reactivation of P-TEFb and latent HIV in primary T cells following either TCR co-stimulation or PKC agonist challenge is independent of PKC but rather involves two complementary signaling arms of the TCR cascade, namely, RasGRP1-Ras-Raf-MEK-ERK1/2 and PI3K-mTORC2-AKT-mTORC1.

**Author Summary:** Dissecting the cellular pathways through which HIV emerges from latency is a key step in the development of therapeutically viable approaches for latency reversal and eventual clearance of persistent HIV in infected individuals. The essential host transcription elongation factor P-TEFb, a heterodimer of CDK9 kinase and a regulatory cyclin T subunit, is a critical mediator of the trans-activation of latent HIV. Availability of P-TEFb for proviral transcription is highly limited due to a posttranscriptional restriction in cyclin T1 expression and dephosphorylation of CDK9 on its activation loop. Using a well-characterized primary T-cell model of HIV latency alongside healthy donor memory CD4^+^ T cells, we have now defined the signaling pathways that are essential for the generation of transcriptionally active P-TEFb and, consequently, proviral reactivation. Crucial among these findings is the demonstration that protein kinase C (PKC) agonists signal through off-target activation of RasGRP1-Ras-Raf-MEK-ERK1/2 to effect the reactivation of both P-TEFb and proviral HIV. Understanding these pathways should lead to the discovery of novel highly selective activators of P-TEFb to improve the efficiency of HIV reactivation in the memory T-cell population of virally suppressed individuals.

## Introduction

Comprehensively understanding the host mechanisms that permit HIV to emerge from transcriptional latency is crucial for the development of safe pharmacological strategies to eliminate persistent viral reservoirs that are primarily found within memory CD4^+^ T cells of infected individuals [1–5]. The switch between HIV latency and productive transcription is regulated by an auto-feedback mechanism initiated by the viral trans-activator protein Tat, which functions to recruit the essential host transcription elongation factor P-TEFb along with the super elongation complex (SEC) to proviral HIV [6–9]. P-TEFb is a heterodimer of CDK9 serine/threonine kinase and one of three cyclin T regulatory subunits. However, cyclin T1 (CycT1) is the only cyclin capable of interacting with HIV Tat [7, 10, 11].

Because of Tat’s auto-feedback mechanism, the transcriptional inactivation of HIV to achieve latency occurs in the absence of Tat expression as acutely infected T cells revert to a resting memory phenotype with the consequent restriction of the nuclear activity of P-TEFb [12, 13]. Subsequently, this results in formation of epigenetic structures at the fixed nucleosome around the viral promoter region (Nuc-1) which hinder the efficient recruitment of RNA polymerase II (RNAP II). The functional importance of these repressive epigenetic modifications is clearly evidenced by the observations that small molecule compounds that inhibit histone deacetylation or methylation can reactivate transcriptionally silenced HIV in cell line and primary cell models of HIV latency [14, 15]. However, even though histone deacetylase inhibitors are thus far the only class of latency reversal agents (LRAs) to achieve clinical proof-of-concept, they have been found to be marginally effective at activating latent proviruses in both primary cell models of HIV latency and patient-derived CD4^+^ T cells [16–22]. In a recent clinical study of the HDACi romidepsin we found that poor HIV reactivation was associated with its limited ability to reactivate P-TEFb and transcription initiation factors [23].

Maintenance of a transcriptionally inactive state in resting memory CD4^+^ T cells, which engenders the persistence of proviral HIV in latently infected cells, is associated with a tight restriction imposed on P-TEFb activity [24, 25]. T-cell receptor (TCR) activation of primary CD4^+^ T cells potently provides the intracellular signals that are necessary to efficiently stimulate processive HIV transcription by reversing the epigenetic silencing at the proviral promoter, inducing the assembly of P-TEFb and permitting nuclear entry of transcription initiation factors [12, 13]. Therefore, the development of more efficient latency reversal strategies will likely require a combination of agents that mimic the effects of TCR signaling.

Physiological TCR signaling in CD4^+^ T cells occurs in the context of an antigen presenting cell (APC) that has on its surface pathogen-derived peptides bound to MHC Class II transmembrane proteins and involves simultaneous activation of multiple signaling pathways. For maximal T-cell activation to be enabled, binding of the peptide-MHC complex to the TCR is accompanied by a costimulatory engagement by the APC that usually involves a pairwise linkage between B7 and its T-cell counterpart CD28 [26]. TCR co-stimulation immediately induces a tyrosine phosphorylation cascade at the cell membrane that leads to the activation of the lipid metabolizing enzymes phospholipase C-γ (PLC-γ) and phosphoinositide-3-kinase (PI3K) [27]. In turn, PLC-γ hydrolyzes phosphatidylinositol-4,5-bisphosphate (PIP_2_) to generate the second messengers inositol-1,4,5-triphosphate (IP_3_), that mobilizes calcium release into the cytoplasm, and diacylglycerol (DAG), which directly contributes to the activation of a number of C1 domain-containing intracellular signaling proteins including protein kinase C (PKC) enzymes and the Ras guanine nucleotide exchange factor RasGRP [28–30]. Class I PI3K enzymes are directly bound to CD28 at the T-cell plasma membrane via a well-characterized YMNM motif on the cytoplasmic domain of the costimulatory receptor [31–33]. Upon their activation, PI3Ks act to phosphorylate PIP_2_ to form phosphatidylinositol-3,4,5-trisphosphate (PIP_3_), a lipid resistant to PLC-γ hydrolysis, that serves to anchor a wide range of signaling proteins through their pleckstrin homology (PH) domains [34].

Biochemical experiments carried out using both Jurkat T-cell and primary cell models of HIV latency [35–39], showed that the activation of latently infected resting primary CD4^+^ T cells upon TCR engagement triggers the rapid nuclear induction of transcription initiation factors such as NFAT, AP-1 and NF-κB. Once in the nucleus these factors bind the proviral promoter and initiate transcription by recruiting histone modifying and chromatin remodeling enzymes whose activity on chromatin at or around the transcription start site allows for formation of the preinitiation complex (PIC) and promoter recruitment of RNAP II.

In contrast to Jurkat T cells, primary resting CD4^+^ T cells are also highly deficient in the expression of the CycT1 subunit of P-TEFb while the CDK9 kinase subunit is sequestered in an inactive form in the cytoplasm bound to the kinase-specific chaperone complex Hsp90/Cdc37 [24, 25]. P-TEFb has been found to be restricted in resting T cells by a combination of microRNA-mediated repression of CycT1 protein synthesis and dephosphorylation of CDK9 on its regulatory activation loop (T-loop) at the highly conserved residue Thr186 [40, 41]. Activation of resting T cells stimulates a rapid posttranscriptional synthesis of CycT1, which is tightly coupled with Thr186 phosphorylation of the CDK9 subunit (pThr186 CDK9), a posttranslational modification that is critical for the stable heterodimeric assembly of P-TEFb [25]. Once P-TEFb is assembled it is incorporated into 7SK snRNP, a nuclear regulatory ribonucleoprotein complex that serves to inhibit its kinase activity and sequester the enzyme in nucleoplasmic substructures called speckles found in close proximity to genomic sites of active transcription [42–45]. Recruitment of P-TEFb from 7SK snRNP by Tat and their cooperative binding to a nascently synthesized RNA hairpin called TAR puts the enzyme in close proximity to the negative elongation factors NELF and DSIF, and promoter-proximally paused RNAP II. CDK9 then acts to phosphorylate the E subunit of the repressive NELF complex at multiple serine residues which causes NELF to dissociate from TAR [46–48]. CDK9 also extensively phosphorylates the CTD of RNAP II mainly at Ser2 residues of the heptad repeats Y-S-P-T-S-P-S and C-terminal repeats (G-S-Q/R-T-P) of the hSpt5 subunit of DSIF at Thr4 residues [49–51]. The overall effect of these phospho-modifications by P-TEFb is to remove the blocks to elongation imposed by NELF and DSIF and to stimulate efficient elongation and co-transcriptional processing of proviral transcripts.

7SK snRNP comprises a molecule of 7SK snRNA that is bound at its 5’ and 3’ ends by the capping enzyme MEPCE and the La-related protein LARP7, respectively [52, 53]. The three-dimensional folding conformation of 7SK snRNA establishes a molecular scaffold that cooperatively binds two molecules of P-TEFb to a homodimer of the inhibitory protein HEXIM1 [54, 55]. Incorporation of P-TEFb into 7SK snRNP requires both Thr186 phosphorylation of CDK9 and the expression of HEXIM1 [25, 56–58], both of which have restricted expression in resting memory CD4^+^ T cells [24]. In cells that are actively dividing, it’s estimated that at least half of all cellular P-TEFb is found in 7SK snRNP [59], clearly denoting an important role this ribonucleoprotein complex plays in regulating global transcription elongation. In a proteomics study designed to identify posttranslational modifications on HEXIM1 that regulate the sequestration of P-TEFb by 7SK snRNP, we found that HEXIM1-associated 7SK snRNP can also serve to retain P-TEFb in the nucleus most likely to provide an exchangeable pool of the enzyme that can be readily availed to stimulate gene transcription [57]. An additional CDK9 T-loop phospho-modification occurs at a conserved residue Ser175 (pSer175 CDK9) which we recently demonstrated is due to CDK7 activity in both Jurkat and primary CD4^+^ T cells [25]. pSer175 CDK9 is only found on a subset of P-TEFb that is transcriptionally active (ie. P-TEFb dissociated from 7SK snRNP) and is functionally important in being coopted by Tat to enhance Tat’s interaction with P-TEFb in order to outcompete BRD4 [25, 60], a bromodomain-containing protein that is considered to be a major recruiter of P-TEFb to cellular genes [59, 61].

Our current study focuses on defining signaling pathways and T-cell factors that are responsible for regulating proviral reactivation in memory T cells, with a specific focus on identifying the pathways that are essential for the generation of transcriptionally active P-TEFb. PKC agonists are well known and highly potent HIV latency reversing agents, but although progress has been made with development of novel synthetic agonists, they have had limited clinical applications due to concerns about their side effects and cytotoxicity [62, 63]. Using a well-characterized and highly reproducible primary QUECEL (quiescent effector cell latency) model of HIV latency [64], alongside healthy donor memory CD4^+^ T cells, we found that diacylglycerol-mimicking PKC agonists can stimulate active P-TEFb expression with minimal cytotoxicity in the absence of co-treatment with a calcium ionophore. Unexpectedly, PKC agonists induce P-TEFb via off-target activation of RasGRP1-Ras-Raf-MEK-ERK1/2. P-TEFb induction after TCR co-stimulation is further regulated by the PI3K-mTORC2-AKT-mTORC1 pathway. Combined inhibition of the PI3K-mTORC2-AKT-mTORC1 pathway and the ERK1/2 activator MEK prior to TCR co-stimulation completely abrogated active P-TEFb expression and substantially suppressed latent HIV reactivation. Thus, reactivation of P-TEFb in primary T cells in response to either TCR co-stimulation or PKC agonist challenge is largely independent of PKC suggesting novel opportunities to develop effective latency reversing strategies.

## Results

### Posttranscriptional induction of P-TEFb expression in primary T cells is required for latent proviral reactivation

Resting CD4^+^ T cells serve as the primary reservoir for latent HIV and maintain a cellular environment that is non-permissive for HIV transcription. Unstimulated memory CD4^+^ T cells isolated from healthy donor blood (**S1A Fig**), the majority of which possess a central memory phenotype (**S1B Fig**), are largely in a resting or quiescent state. Characteristically, the resting cells lack markers of T-cell activation (CD25 and CD69), cell proliferation (Ki67 and cyclin D3) and critical transcription factors such as NF-κB and P-TEFb (**S2 and S3 Figs**).

As shown in **Fig 1A**, P-TEFb activity is highly restricted in resting memory CD4^+^ T cells due to a posttranscriptional block in CycT1 translation and sequestration of activation-loop-dephosphorylated CDK9 in the cytoplasm by Hsp90/Cdc37 [25, 40, 41]. We have developed sensitive flow cytometry assays to monitor CycT1 expression and phosphorylation of CDK9 on its activation loop (T-loop) after stimulation of memory CD4^+^ T cells (**Figs 1B and 1C**) [25, 60]. Using this assay with a range of known latency reversing agents revealed that T-cell receptor (TCR) co-stimulation with anti-CD3 and anti-CD28 coated beads is the most effective at eliciting induction of CycT1 expression and T-loop phosphorylation of CDK9 in memory CD4^+^ T cells (**Fig 1D**). Activation of memory T cells through the TCR results in a rapid synthesis of CycT1 protein (**Fig 1B and S3 Fig**) and, just as rapid, the localization of nascent P-TEFb into the nucleus (**Fig 2A**). This process is coincident with the assembly of the P-TEFb regulatory complex 7SK snRNP (**Fig 2B**). The phorbol ester ingenol, prostratin and PMA were also effective inducers of P-TEFb, but not surprisingly, almost no P-TEFb was induced by the relatively ineffective HDACi SAHA or by stimulation of cells with anti-CD3 alone, ionomycin or TNF-α (**Fig 1D**).

**Fig 1.**
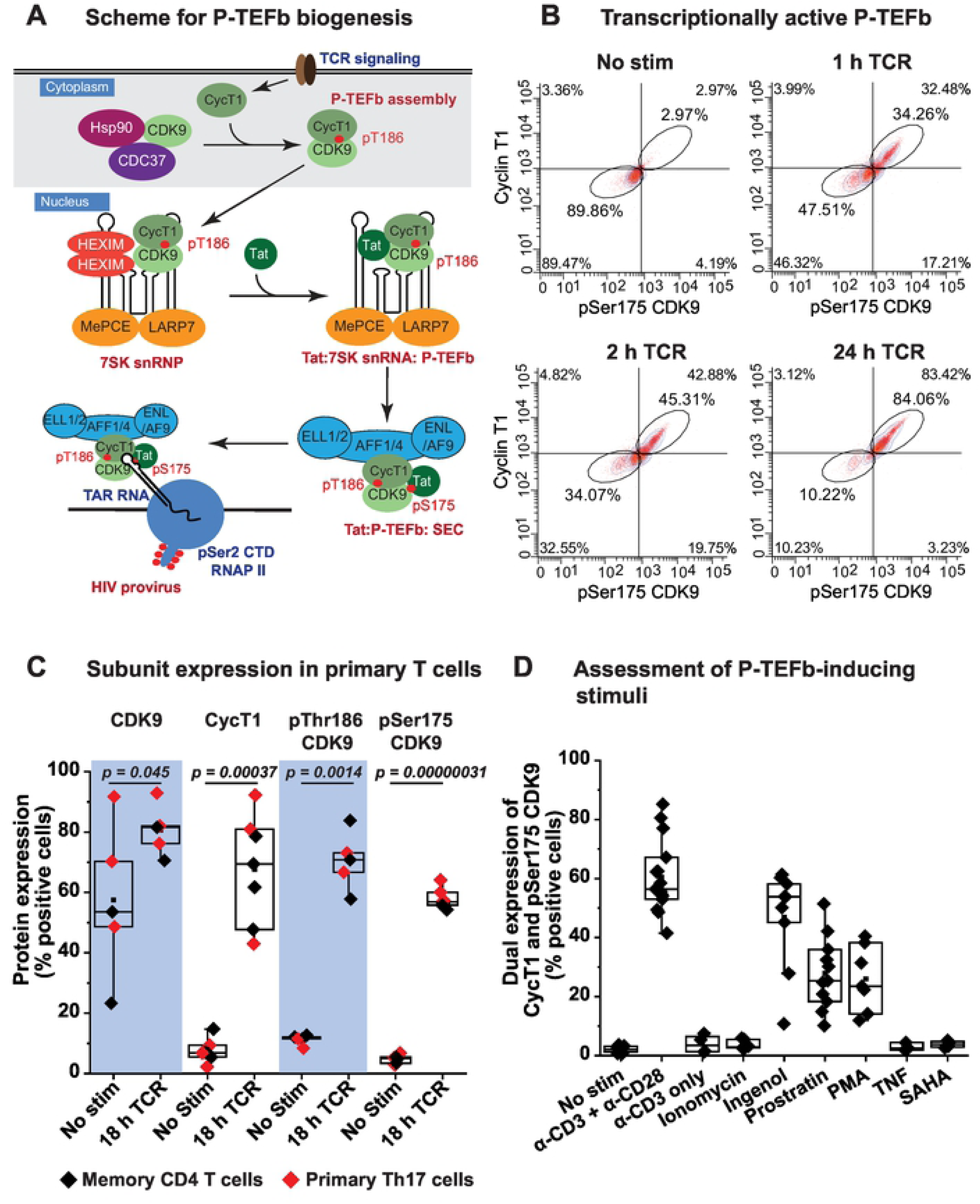
Stimulation of P-TEFb assembly in memory CD4^+^ T cells by TCR co-stimulation and PKC agonists. *(A)* Proposed scheme for the biogenesis of P-TEFb in memory CD4^+^ T cells and its exchange from 7SK snRNP to assemble the super elongation complex (SEC) on proviral HIV. Posttranscriptional synthesis of cyclin T1 (CycT1) initiated immediately upon TCR co-stimulation serves as a trigger for the heterodimeric assembly of P-TEFb and T-loop phosphorylation of CDK9 kinase at Thr186. Assembled P-TEFb is translocated into the nucleus where it is associated with HEXIM1 and 7SK snRNP to form 7SK snRNP. Exchange of P-TEFb from 7SK snRNP is facilitated by the displacement of HEXIM1 by Tat and also T-loop phosphorylation of CDK9 at Ser175 leading to formation of the super elongation complex (SEC) containing transcriptionally active P-TEFb. This complex is then loaded onto TAR RNA to stimulate proviral transcription elongation. Signaling pathways responsible for the biogenesis of P-TEFb leading up to formation of transcriptionally active P-TEFb will be examined in this study. *(B)* Assessment of active P-TEFb expression by immunofluorescence flow cytometry as measured by monitoring the co-expression of cyclinT1 and pSer175 CDK9. *(C)* Flow cytometry analysis of the intracellular fluorescence immunostaining of total CDK9, cyclin T1 (CycT1), pThr186 CDK9 and pSer175 CDK9 from at least five different experiments performed using either primary Th17 (red) or memory T cells (black) that were activated or not for 18 h through the TCR with anti-CD3/anti-CD28 Dynabeads. *(D)* Assessment of stimuli capable of inducing P-TEFb expression in memory CD4^+^ T cells. The graph shows immunofluorescence flow cytometry measurements of the coordinate induction of CycT1 and pSer175 CDK9 expression in memory T cells derived from at least four different healthy donors. Cells were subjected to 24 h treatment with the indicated stimuli prior to immunofluorescence staining for CycT1 and pSer175 CDK9.

**Fig 2.**
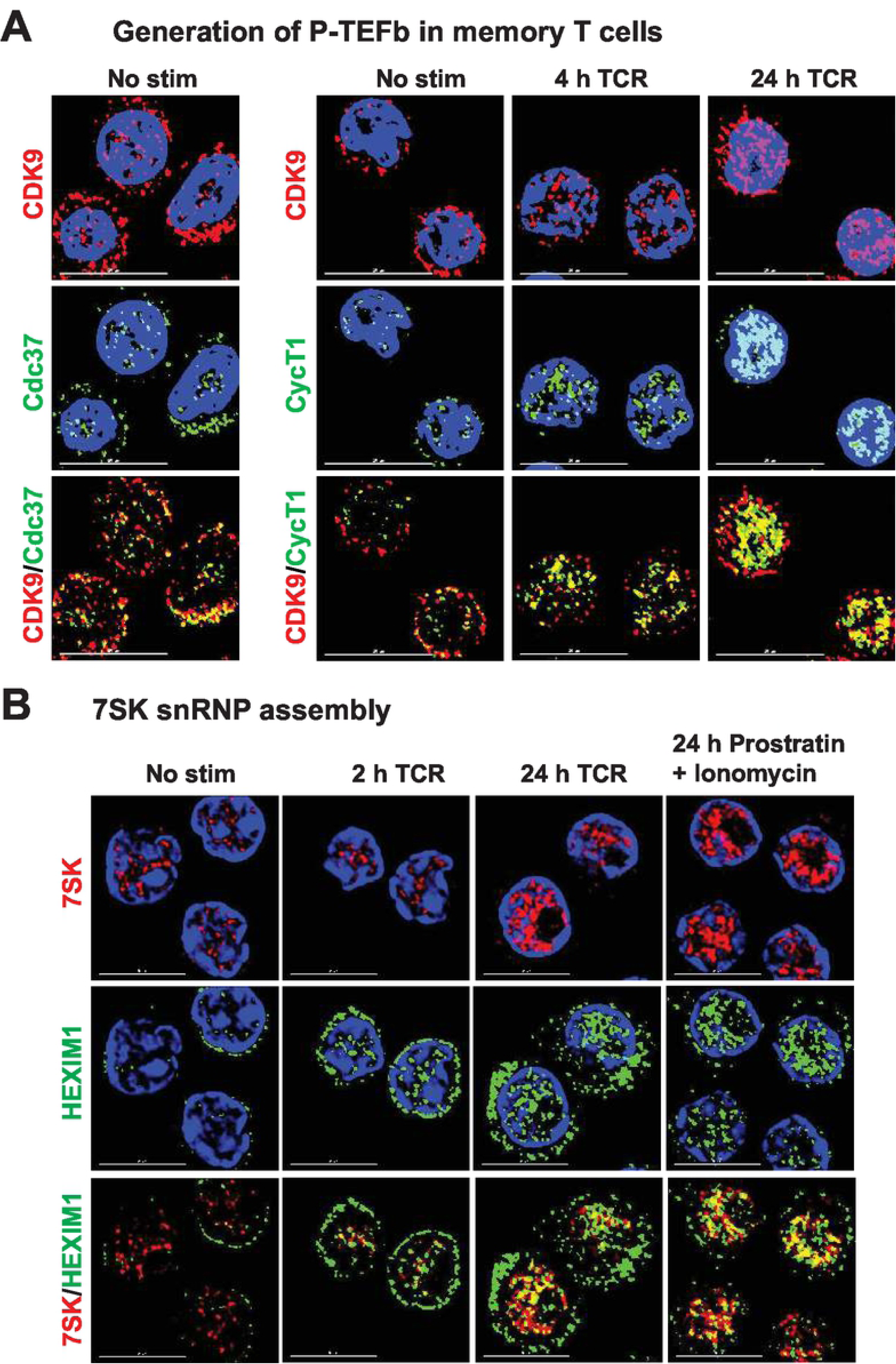
Imaging the nuclear assembly of P-TEFb and 7SK snRNP in activated memory CD4^+^ T cells. *(A)* Subcellular expression of P-TEFb subunits in memory CD4^+^ T cells before and after T-cell receptor (TCR) activation. Cells were stimulated or not with a soluble cocktail of anti-CD3/anti-CD28 antibodies for 4 or 24 h prior to dual immunostaining for CDK9 and CycT1. Images were captured at 100X using a deconvolution microscope. Scale bar: 10 µm. *(B)* Visualization of the expression and subcellular localization of 7SK snRNA and HEXIM1 in memory CD4^+^ T cells. These imaging experiments were carried out by combining RNA FISH with immunofluorescence staining of unstimulated memory T cells, cells activated through the TCR for 2 or 24 h with anti-CD3/anti-CD28 Dynabeads, or cells challenged with a combination of the PKC agonist prostratin and the calcium ionophore ionomycin. Images were captured at 100X by deconvolution microscopy. Scale bar: 5 µm.

As shown in **Fig 1C** the expression of P-TEFb subunits and the T-loop phosphorylated forms of CDK9 in both healthy donor memory CD4^+^ T cells and resting polarized primary Th17 cells obtained using the QUECEL protocol (**S4A Fig**) before and after TCR activation are indistinguishable. We have extensively used the highly reproducible QUECEL model to study HIV latency in primary CD4+ T cells [15, 64]. In the unstimulated condition, both memory CD4^+^ T cells and resting primary Th17 cells exhibit low expression of CycT1 and the T-loop phosphorylated forms of CDK9 (pSer175 and pThr186) (**Fig 1C and S2, S3, S4B Figs**). While TCR activation of both cell types over an overnight period resulted in a modest elevation of CDK9 protein (by an average of 1.4-fold), it also led to a substantial and proportional increase in the percentage of cells that express CycT1, pSer175 CDK9 and pThr186 CDK9 (by an average of 9-fold, 12.2-fold, and 6.3-fold, respectively) (**Fig 1C**). Generation of CycT1 and the transcriptionally active marker of P-TEFb, pSer175 CDK9, were observed to be quite rapid, reproducibly occurring within 2 h of TCR co-stimulation and concomitant with formation of the P-TEFb-phosphorylated form of RNAP II (pSer2 RNAP II CTD) (**Fig 1B and S3, S5A Figs**). These results clearly indicate that the dual expression profiling of CycT1 and pSer175 CDK9 can serve as a dependable measure of catalytically active and gene-associated P-TEFb in primary T cells. Therefore, in the current study induction of transcriptionally active P-TEFb was assessed as the coordinate expression of CycT1 and pSer175 CDK9.

The absolute requirement for P-TEFb induction to support emergence of HIV from latency in primary CD4^+^ T cells was demonstrated by the observation that a relatively selective CDK9 kinase inhibitor flavopiridol (FVP) repressed proviral reactivation in latently infected primary Th17 cells by an average of 2.5-fold (**S5B Fig**). Therefore, HIV reactivation in latently infected primary T cells is clearly preceded by and dependent on CycT1 posttranscriptional synthesis and the formation of transcriptionally active P-TEFb in the nucleus in response to intracellular signals emanating from TCR co-stimulation.

### Regulation of 7SK snRNP and super elongation complex factors in primary T cells

Efficient stimulation of proviral HIV transcription elongation by P-TEFb is regulated by 7SK snRNP and facilitated by the super elongation complex (SEC) (**Fig 1A**) [8, 9, 42, 43]. We therefore examined the expression profiles of 7SK snRNP and SEC components in primary CD4^+^ T cells before and after activation. Western blotting of resting and TCR-activated Th17 cells demonstrated notable restrictions in protein expression of not just CycT1 but also HEXIM1 and the SEC components AFF1 and ELL2 during quiescence (**S6A Fig**). Transcriptome analysis (bulk RNA-seq) of primary CD4^+^ T cell types that had been polarized into the four major T-cell subsets (Th1, Th2, Th17, and Treg) [64] was also conducted to provide an additional measure of the differential expression of transcripts belonging to P-TEFb subunits (CDK9, CycT1 and CycT2), 7SK snRNP components (HEXIM1, HEXIM2, LARP7 and MEPCE) and SEC proteins (AFF1, AFF2, ELL, ELL2 and ENL). By including in our analysis a publicly available bulk RNA-seq dataset generated by Zhao et al. using unstimulated and TCR-activated memory CD4^+^ T cells [65] (SRA accession SRP026389) (Extended data shown in **S6B-S6D Figs**), we were able to demonstrate that each of these *in vitro* polarized T-cell subsets can faithfully recapitulate the phenotypic characteristics of memory CD4^+^ T cells in maintaining the pattern of expression of these factors (**Figs 3A-3C**). Most notably, the high-throughput sequencing studies demonstrated that there was little change in CycT1 mRNA levels over a 24-h period of TCR co-stimulation (**Fig 3A and S6B Fig**), consistent with the prevailing idea that this P-TEFb subunit is post-transcriptionally regulated. Also noteworthy is the observation that HEXIM2 mRNA was very low in resting primary CD4^+^ T cells (~6 transcripts per million) and was further slightly reduced following TCR co-stimulation (~5 transcripts per million) (**Fig 3B**). By contrast, HEXIM1 was found to be expressed at approximately 4-fold and 7-fold higher levels compared to HEXIM2 in unstimulated and 24-h TCR-activated primary CD4^+^ T cells, respectively (**Fig 3B**). This finding suggests that HEXIM1 is the major isoform in CD4^+^ T cells and therefore the more probable regulatory partner for P-TEFb within 7SK snRNP. Consistent with our immunoblotting analysis (**S6A Fig**) and previous findings that ELL2 is tightly regulated and therefore an important limiting factor for SEC formation [66, 67], we observed that ELL2 mRNA is minimal in resting CD4^+^ T cells and highly inducible in response to TCR activation (**Fig 3C and S6C Fig**). Single-cell RNA-seq (scRNA-seq) that were carried out using both resting and TCR-activated memory CD4^+^ T cells also supported the findings from the bulk RNA-seq analysis in showing a restriction of ELL2 in resting T cells and a predominance of HEXIM1 over HEXIM2 (**S7A and S7B Figs**). Thus, these overall results suggest that the posttranscriptional restriction in CycT1 and repression of the P-TEFb-associated factors HEXIM1, AFF1 and ELL2 may need to be overcome in resting CD4^+^ T cells in order to achieve a robust reactivation of HIV from latency.

**Fig 3.**
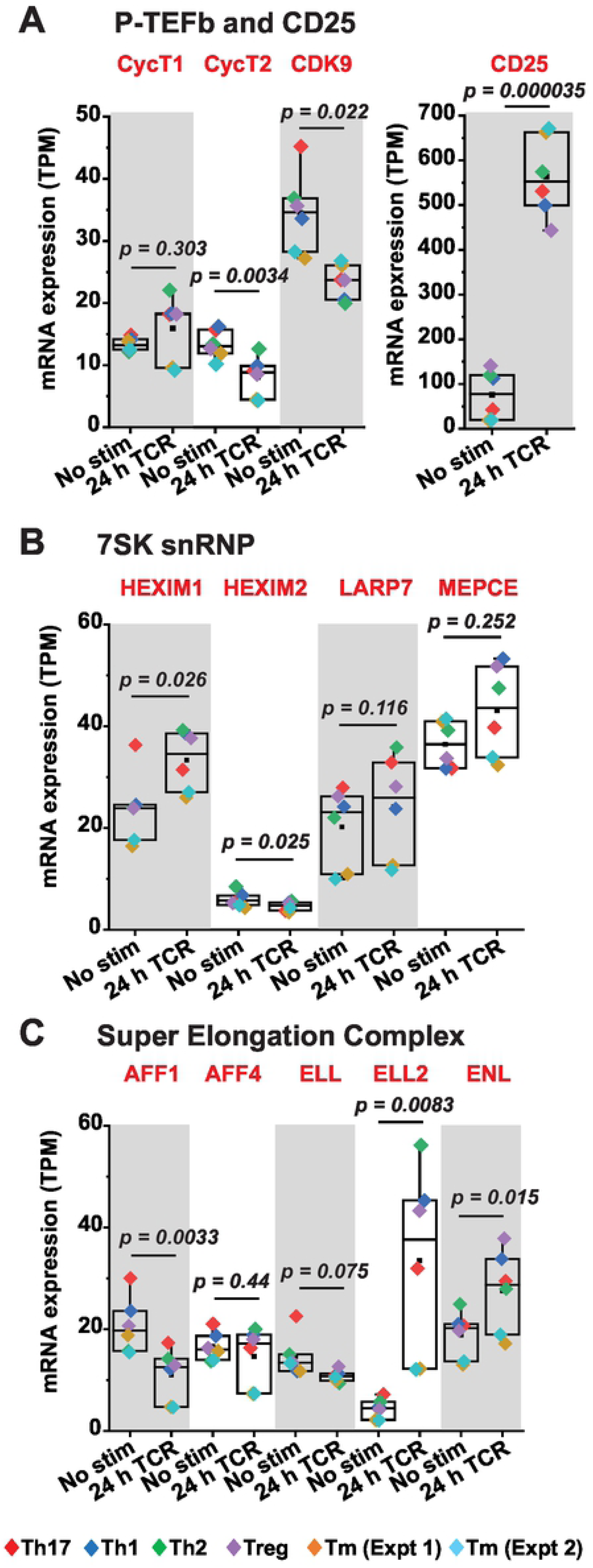
Transcriptome analysis of P-TEFb, 7SK snRNP and SEC expression in primary QUECEL subsets and memory CD4^+^ T cells. Quiescent CD4**^+^** T cells that had been polarized into Th1, Th2, Treg and Th17 subsets using the QUECEL procedure were activated or not for 24 h with anti-CD3/anti-CD28 Dynabeads. Bulk RNA-seq datasets obtained using these cells were analyzed to examine the expression of P-TEFb subunits *(A)*, 7SK snRNP components *(B)*, and factors belonging to the super elongation complex *(C)*. Transcripts per million (TPM) values were used to evaluate the relative abundance of transcripts under resting and activated conditions. In *A*, the expression of CD25 was examined as a positive control. A publicly available bulk RNA-seq dataset of primary human memory CD4^+^ T cells that had been activated or not through TCR co-stimulation (SRA accession SRP026389) with anti-CD3/anti-CD28 coated beads was also analyzed for these factors (Tm Expt 1 and Expt 2). Statistical significance (*p* values) was calculated using a two-tailed Student’s *t* test.

### Reactivation of P-TEFb in primary T cells is not mediated by protein kinase C

Treatment with a variety of protein kinase C (PKC) agonists (ingenol, prostratin or PMA) stimulated the dual expression of CycT1 and pSer175 CDK9 in memory CD4^+^ T cells but at lower mean levels compared to that observed with TCR co-stimulation (**Fig 1D**). Co-treatment of memory T cells with ionomycin and each of the three PKC agonists did not further elevate P-TEFb expression but substantially enhanced formation of both the transcriptionally active form of NF-κB (pSer529 p65) and the P-TEFb-modified pSer2 CTD RNAP II (**Fig 4A**). While treatment with PKC agonists elicited minimal cytotoxicity to memory T cells (relative to unstimulated cells viability was found to be 93%, 91% and 87% for ingenol, prostratin and PMA, respectively), co-treatment with ionomycin resulted in a significant loss of cell viability (35-40% cell death compared to unstimulated cells) (**Fig 4A**). These results suggest that while intracellular calcium mobilization is not essential for enabling P-TEFb biogenesis in memory T cells, it significantly potentiates the efficacy of PKC agonists at generating the transcription elongation competent form of RNAP II albeit with significant cell cytotoxicity. Exposure of polarized primary Th17 cells to ingenol or prostratin was sufficient to stimulate the formation of P-TEFb and resulted in a comparable extent of proviral reactivation in latently infected Th17 cells as TCR co-stimulation (**Fig 4B and S5D Fig**). By comparison, TNF-α and SAHA which are ineffective at generating transcriptionally active P-TEFb in both memory and Th17 cells (**Fig 1D and S4B Fig**), were found to be poor reactivators of proviral HIV in the QUECEL Th17 latency model (**S5C Fig**). Thus, the reactivation of HIV from latency appears to be tightly coupled to the biogenesis of P-TEFb in primary T cells and both of these processes can be elicited upon cellular treatment with PKC agonists in the absence of intracellular calcium mobilization.

**Fig 4.**
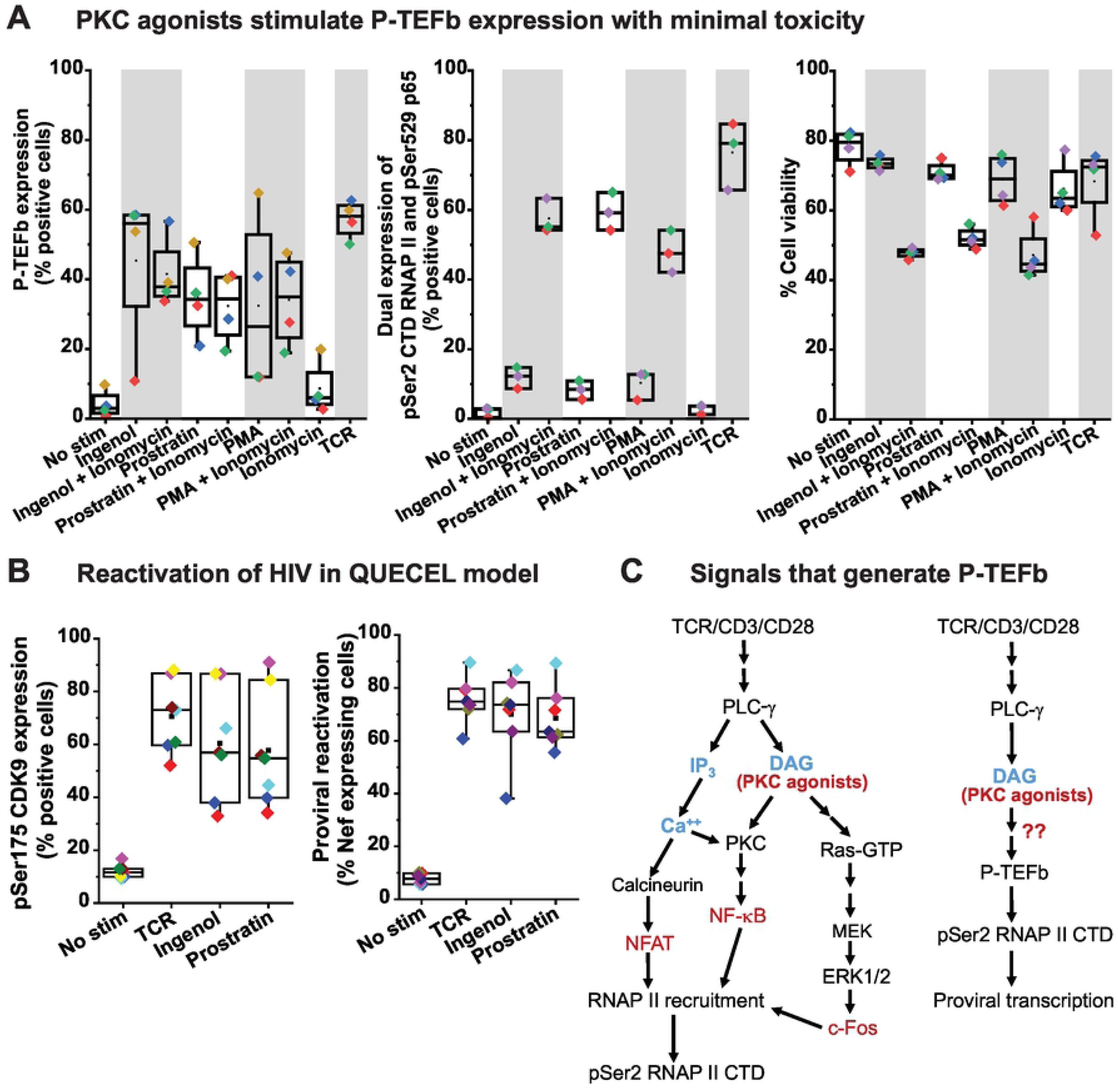
P-TEFb biogenesis in primary T cells does not require intracellular mobilization of calcium. *(A) Left graph*; Cross-comparison of the effectiveness of PKC agonists on their own or combined with ionomycin at inducing active P-TEFb in primary T cells measured by co-staining for CyclinT1 and pSer175 CDK9. *Middle graph*; Cross-comparison of the effectiveness of PKC agonists on their own or combined with ionomycin at coordinately inducing the C-terminal domain Ser2 phosphorylated form of RNA polymerase II (pSer2 RNAP II CTD) and the transcriptionally active form of NF-κB (pSer529 p65). *Right graph*; Measurement of the effect of PKC agonists on their own or combined with ionomycin on cell viability as assessed by flow cytometry following staining with the eFluor450 viability dye. *(B)* Coordinate reactivation of P-TEFb *(left graph)* and latent HIV *(right graph)* upon 24 h treatment of resting Th17 cells with either 50 nM Ingenol or 1 µM Prostratin. The graphs show data generated using cells derived from at least three different donors. *(C)* Proposed schemes for the promoter recruitment of RNA polymerase II (RNAP II) to proviral HIV and its phosphorylation by active P-TEFb on its C-terminal domain (pSer2 RNAP II CTD). Arrows with red question marks indicate a P-TEFb regulatory signaling pathway(s) intended to be defined by the current study.

PKC agonists, commonly referred to as phorbol esters, are structural mimics of endogenously generated diacylglycerol (DAG) and bind the C1 domains of both conventional (α, βI, βII, and γ) and novel (δ, θ, ε, and η) PKC isozymes leading to their membrane recruitment and stimulation of their kinase activities [30]. We therefore examined whether any of these DAG-binding PKC enzymes mediate the expression of CycT1 and/or T-loop phosphorylated CDK9 in primary T cells in response to TCR co-stimulation or phorbol ester treatment (**Fig 4C**). Healthy donor memory CD4^+^ T cells and polarized primary Th17 cells were either activated through the TCR or challenged with phorbol ester in the presence or absence of treatment with a conventional pan-PKC inhibitor (Ro-31-8220), an inhibitor that potently inhibits PKC α, β, γ, and δ isoforms (Gö-6983), or one that selectively targets PKC-θ (sotrastaurin). PKC-θ is considered to be the predominantly functional PKC isoform in T cells and it critically mediates both TCR-induced NF-κB and global T-cell activation [68–70]. Surprisingly, all three PKC inhibitors failed to effectively block either the TCR- or the phorbol ester-mediated inducible expression of CycT1, pSer175 CDK9 and pThr186 CDK9 in both memory CD4^+^ T cells and polarized primary Th17 cells or had only very modest effects in their inhibition (**Figs 5A, 5B and S8 Fig**). Similarly, these inhibitors were ineffective at suppressing the reactivation of latent HIV in primary Th17 cells that was elicited by either TCR co-stimulation or challenge with phorbol ester (**Figs 5B, 5C and S5E Fig**).

**Fig 5.**
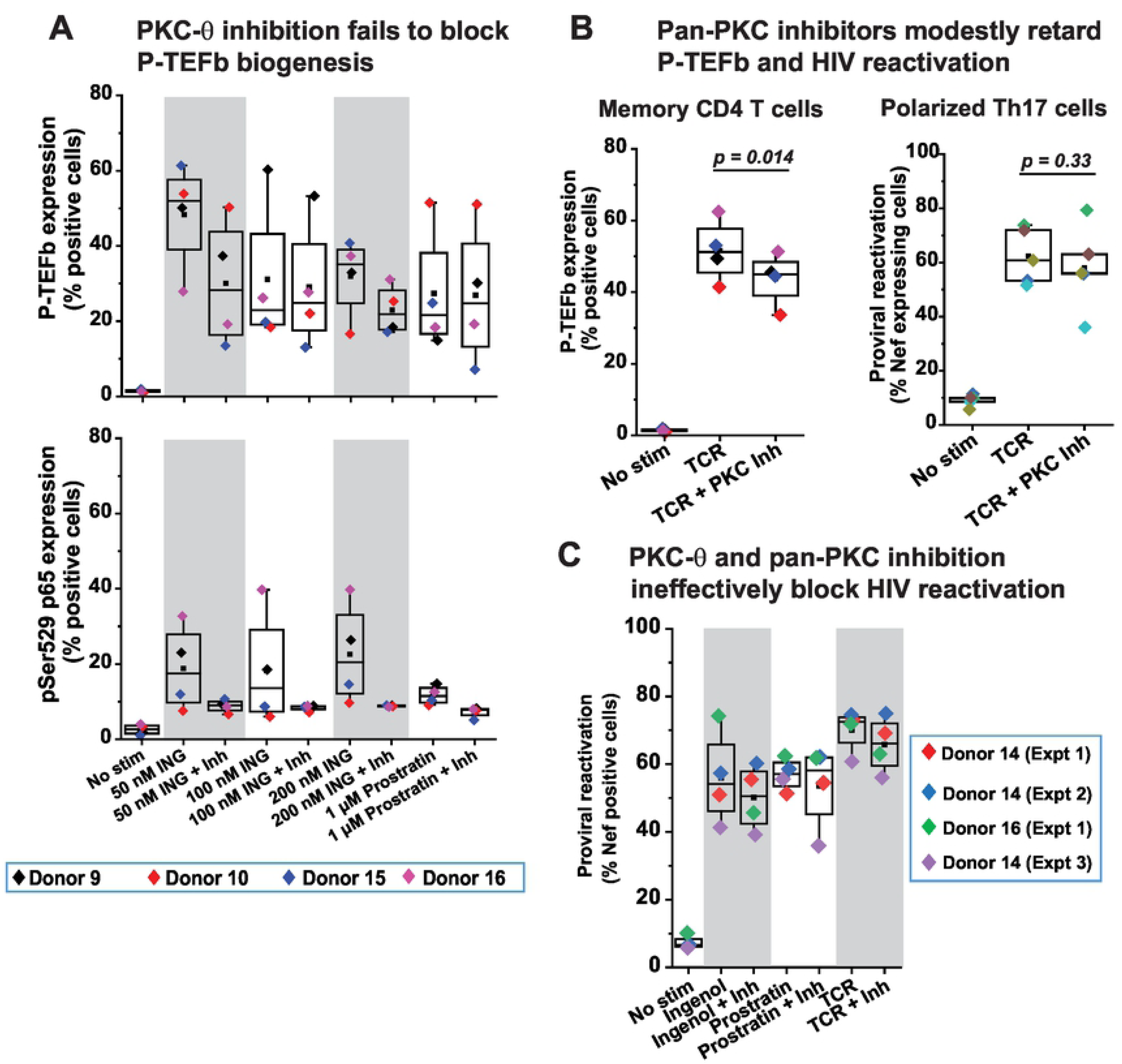
Coordinate reactivation of P-TEFb and latent HIV in primary T cells is largely independent of PKC activity. *(A)* The PKC-θ inhibitor sotrastaurin is ineffective at repressing ingenol- and prostratin-induced expression of P-TEFb but effective at blocking the activating phosphorylation of the p65 subunit of NF-κB at Ser529 in response to these phorbol esters. CD4^+^ memory T cells from four different donors were treated for 24 h with the indicated concentrations of either ingenol or prostratin in the presence or absence of 100 nM sotrastaurin prior to subjecting cells to dual immunofluorescence staining for CycT1 and pSer175 CDK9 *(top graph)* or immunofluorescence staining for pSer529 p65 *(bottom graph)*. *(B)* PKC inhibitors modestly repress TCR-mediated expression of P-TEFb in CD4^+^ memory T cells but are unable to block reactivation of proviral HIV in primary Th17 cells. *Left graph*; Measurement of P-TEFb expression by dual CyclinT1 and pSer175 CDK9 immunofluorescence staining of memory T cells from four different healthy donors that were treated or not with 100 nM Sotrastaurin for 30 min prior to TCR co-stimulation anti-CD3/anti-CD28 Dynabeads for 24 h. *Right graph*; Assessment of the effect of PKC inhibition on proviral reactivation in *ex vivo* HIV-infected primary Th17 cells prepared from three healthy donors. Latently infected cells were treated or not for 30 min with 100 nM Ro-31-8220 in two experiments or with 100 nM Sotrastaurin in three experiments prior to TCR co-stimulation with anti-CD3/anti-CD28 Dynabeads for 24 h. Thereafter, cells were analyzed by flow cytometry following immunostaining using a fluorophore-conjugated antibody towards HIV Nef. Statistical significance (*p* values) was calculated using a two-tailed Student’s *t* test. *(C)* PKC inhibitors are unable to repress the reactivation of latent HIV in primary Th17 cells in response to PKC agonists. Latently infected Th17 cells were treated or not for 30 min with either a combination of Ro-31-8220 and Gö 6983 (Donor 14 (Expt 1) and Donor 14 (Expt 2)) at 100 nM each or 100 nM Sotrastaurin (Donor 16 (Expt 1) and Donor 14 (Expt 3)) prior to TCR co-stimulation or challenge with the PKC agonists shown. Thereafter, cells were immunostained using a fluorophore-conjugated antibody towards HIV Nef and analyzed by flow cytometry.

By contrast, sotrastaurin effectively inhibited the activation of NF-κB in memory T cells in response to phorbol esters as measured by monitoring the formation of pSer529 p65 (**Fig 5A**). In addition to serving as a positive control for the drug’s activity, this observation is consistent with the prevailing idea that PKC-θ is a primary mediator of NF-κB activation in T cells. We were also able to demonstrate that Ro-31-8220 can disrupt the degradation of IκB-α due to TCR activation in memory T cells, thereby delaying the peak mobilization of NF-κB subunits p65 and p50 into the nucleus (**S9A-S9C Fig**). Overall, these results indicate that the coordinate reactivation of P-TEFb and latent HIV in primary T cells in response to either TCR co-stimulation or phorbol ester challenge is largely independent of PKC signaling pathways.

### PKC agonists signal through the MAPK ERK pathway to stimulate P-TEFb expression and reactivate latent HIV in primary T cells

TCR-mobilized diacylglycerol and exogenous phorbol esters can also target several other proteins with C1 domains including RasGRP, a guanine nucleotide exchange factor for Ras in T cells [29]. Membrane-anchored RasGRP1 facilitates the activity of SOS, a second Ras GDP-GTP exchange factor, in a positive feedback mechanism that involves the allosteric activation of SOS by RasGRP-generated Ras-GTP [71]. Exchange of GDP for GTP on Ras by both RasGRP and activated SOS causes the formation of Ras–Raf complexes at the plasma membrane that lead to the activation of the MAPK isoforms ERK1 and ERK2 (ERK1/2) (**Fig 6A**) [72, 73].

**Fig 6.**
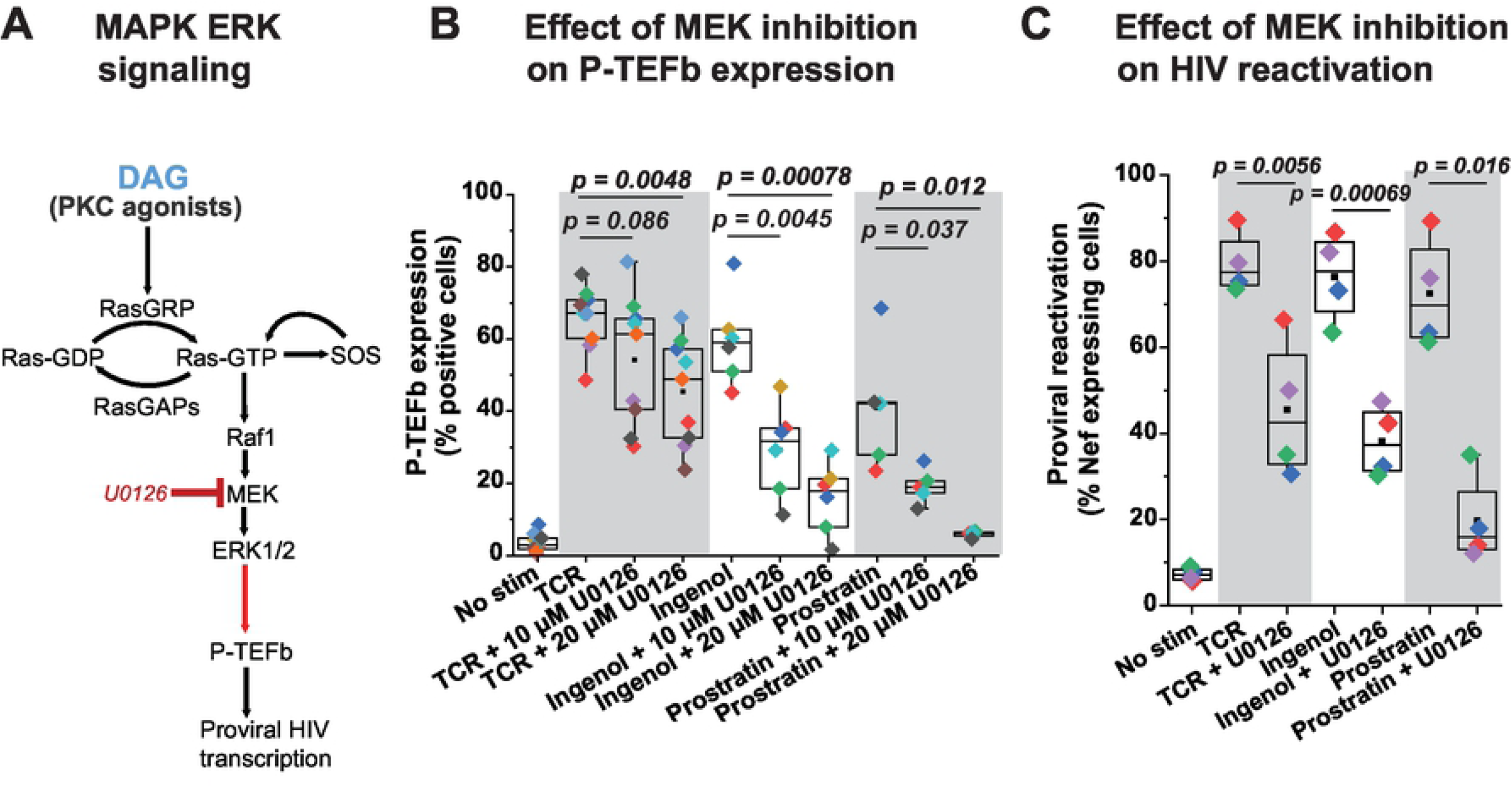
Inducible P-TEFb expression and proviral reactivation in primary T cells are mediated by the MAPK ERK pathway. *(A)* Proposed scheme for the intracellular stimulation of P-TEFb expression in primary T cells by PKC agonists or TCR-generated diacylglycerol (DAG) leading up to the reactivation of HIV from latency. *(B)* The MEK inhibitor U0126 substantially blocks expression of active P-TEFb in primary CD4^+^ T cells in response to treatment with PKC agonists but modestly inhibits P-TEFb expression in response to TCR co-stimulation. Cells were treated or not for 30 min with U0126 at the indicated concentrations prior to TCR co-stimulation with anti-CD3/anti-CD28 Dynabeads or challenge with either 50 nM Ingenol or 1 µM Prostratin for 24 h. Cells were then analyzed by flow cytometry for P-TEFb expression following immunostaining using fluorophore-conjugated antibodies towards CyclinT1 and pSer175 CDK9. *(C)* U0126 substantially retards proviral reactivation in primary Th17 cells in response to TCR co-stimulation or challenge with PKC agonists. Latently infected Th17 cells prepared using naïve CD4^+^ T cells from two different donors were pretreated or not with 20 µM U0126 for 30 min prior to TCR co-stimulation with anti-CD3/anti-CD28 Dynabeads or challenge with 50 nM ingenol or 1 µM prostratin for 24 h. Cells were analyzed by flow cytometry following immunostaining using a fluorophore-conjugated antibody towards HIV Nef. All the *p* values shown in *B* and *C* were calculated using a two-tailed Student’s *t* test.

We were able to confirm that phorbol ester treatment of T cells rapidly stimulates the activating phosphorylation of ERK1/2 (pThr202/pTyr204) concomitant with the inducible synthesis of c-Fos and its localization to the nucleus (**S10A Fig**). Therefore, we examined whether the Ras– Raf–MEK–ERK1/2 pathway (**Fig 6A**) can also mediate the biogenesis of P-TEFb in response to TCR or phorbol ester challenge and if it could be a primary pathway for the reactivation of latent HIV in primary CD4^+^ T cells.

Treatment of Th17 and CD4^+^ memory T cells with the MEK inhibitor U0126, which effectively blocks the activating phosphorylation of ERK1/2 (**S10A and S10B Figs**), resulted in a modest inhibition of TCR-induced P-TEFb expression (by an average of 1.5-fold) (**Fig 6B**). By contrast, U0126 treatment much more effectively inhibited the expression of P-TEFb in both Th17 and memory T cells stimulated by either ingenol or prostratin (by an average of 3.7-fold and 7-fold, respectively) (**Figs 6B and S11-S13 Figs**). Consistent with this observation, U0126 also substantially repressed proviral reactivation in HIV-infected Th17 cells that was induced by ingenol and prostratin (by an average of 2-fold and 3.7-fold, respectively) (**Fig 6C and S14 Fig**).

Although U0126 had a modest and variable repressive effect on TCR-induced P-TEFb expression, there was a stronger repressive effect on TCR-induced proviral reactivation in HIV-infected Th17 cells (1.75-fold repression of HIV compared to the 1.5-fold repression observed on P-TEFb) (**Fig 6C and S14 Fig**). To test the hypothesis that TCR co-stimulation may be exerting effects on the initiation of proviral transcription by generating the activator protein 1 (AP-1) transcription factor complex of c-Fos and c-Jun through a crosstalk between the ERK1/2 and JNK MAPK pathways (**S15A Fig**), we examined whether inhibiting JNK either individually or in combination with MEK had any significant effects on proviral reactivation as well as P-TEFb expression. Inhibition of JNK signaling with SP600125 caused a 1.3-fold reduction in TCR-induced proviral reactivation compared to a 1.7-fold reduction observed with U0126 treatment (**S15B Fig**). Treatment of latently infected Th17 cells with both inhibitors failed to elicit additional suppression of TCR-induced proviral gene expression beyond that observed with U0126 treatment (**S16 Fig**). Similarly, inhibition of JNK did not further enhance the modest repression of TCR-induced P-TEFb expression observed upon MEK inhibition (**S17 Fig**). Moreover, and not surprisingly, inhibition of JNK had a very modest suppressive effect on ingenol- and prostratin-induced proviral and P-TEFb expression (**S16 and S17 Figs**) clearly indicating that these phorbol esters do not signal through JNK in eliciting their stimulatory effects. By contrast, a Raf inhibitor AZ 628 which selectively and potently inhibits Raf1 and B-Raf kinase activities (with IC_50_ values of 29 and 105 nM, respectively) was effective at suppressing P-TEFb induction in response to ingenol and prostratin but had no inhibitory effect on TCR-induced P-TEFb expression (**S18 Fig**).

In summary, our results clearly suggest that MEKK1-MKK4/7-JNK-c-Jun is unlikely to be an essential MAPK pathway for regulating the formation of P-TEFb or the reactivation of latent HIV in CD4^+^ T cells. Instead, our results strongly support the idea that the Raf-MEK-ERK1/2 MAPK pathway is a primary mediator of the expression of active P-TEFb in memory T cells upon challenge with DAG-mimicking phorbol esters and that this signaling pathway is also essential for the reactivation of latent HIV by these candidate LRAs.

### RasGRP1 is the predominantly expressed DAG-dependent RasGRP isoform in primary CD4^+^ T cells

ERK1/2 activation by TCR co-stimulation is dependent on stimulation of Ras by its membrane-associated guanine nucleotide exchange factors RasGRP and SOS [74]. The Ras GDP-GTP exchange activity of SOS in T cells has been found to be allosterically dependent upon initial activation of Ras by RasGRP1 [71]. Therefore, we hypothesized that RasGRP1 has a central role in mediating the reactivation of P-TEFb and HIV in primary T cells through the ERK1/2 pathway upon challenge with PKC agonists.

The guanine nucleotide exchange activity of three of the four known isoforms of RasGRP, namely, RasGRP1, RasGRP3 and RasGRP4 is known to be tightly regulated by the high affinity binding of membrane-associated DAG or phorbol esters to the C1 domains of these proteins [29]. Upon cellular treatment with DAG-mimicking phorbol esters, RasGRP1, RasGRP3 and RasGRP4 rapidly translocate to cellular membranes where they contribute to the activation of membrane-bound Ras [75–77]. By contrast, the C1 domain of RasGRP2 has a very poor binding affinity for DAG [78, 79], and consistent with this observation, the protein fails to translocate to cellular membranes in response to treatment with DAG analogs [80].

We therefore examined whether treatment of memory CD4^+^ T cells with PKC agonists can trigger the recruitment of RasGRP1 to the plasma membrane. High magnification immunofluorescence microscopy revealed a rapid (within 2 h) translocation of RasGRP1 to the membrane following stimulation of memory T cells with ingenol, prostratin or PMA (**Fig 7A**). The degree of colocalization of RasGRP1 with the plasma membrane-resident protein Ezrin in cells treated with each of these three PKC agonists was comparable to that observed with TCR-activated cells (**Fig 7B**).

**Fig 7.**
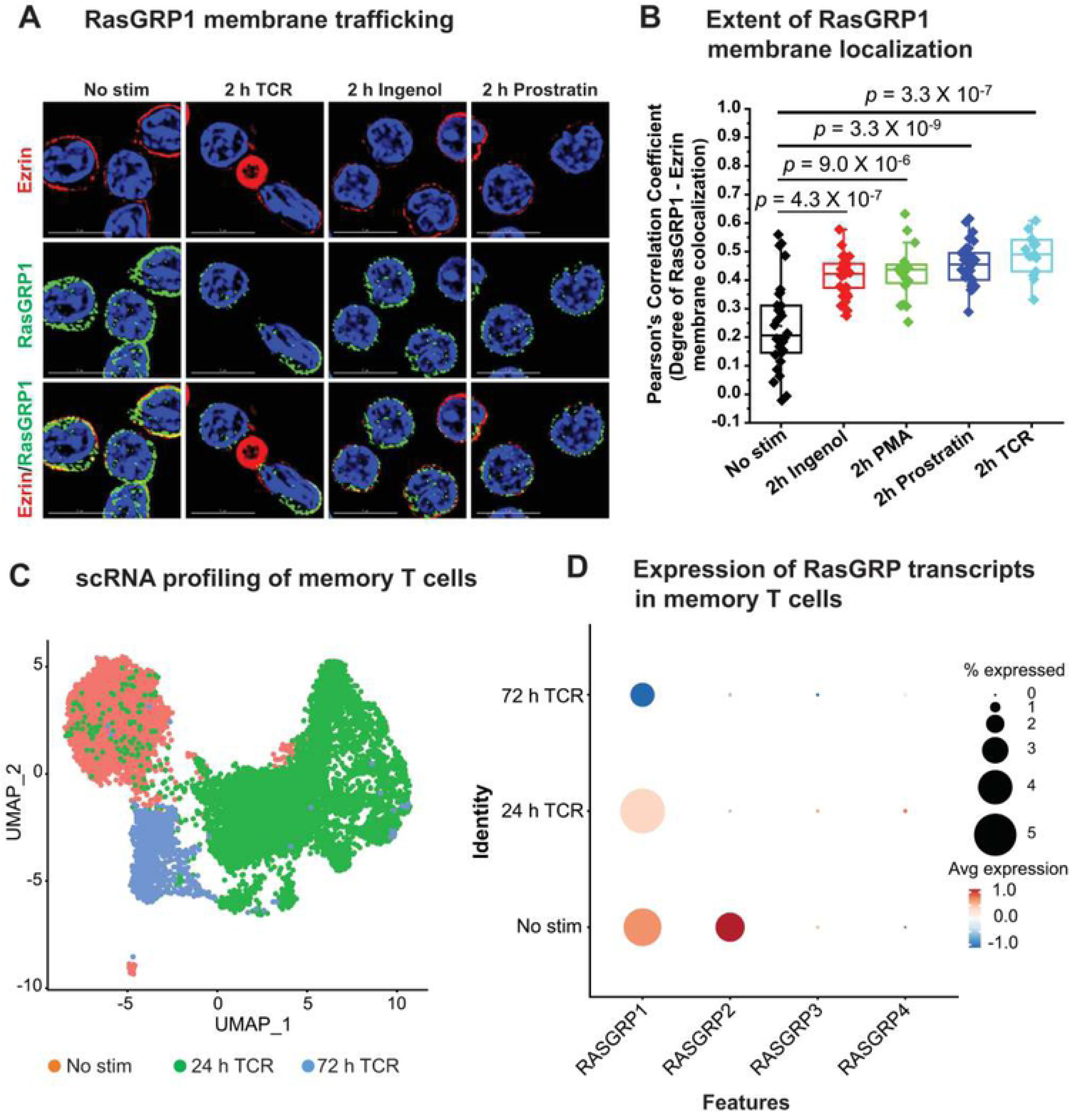
Memory CD4^+^ T cells predominantly express the RasGRP1 isoform which rapidly localizes to the plasma membrane upon phorbol ester stimulation. *(A)* Immunofluorescence microscopy of healthy donor memory CD4^+^ T cells with or without a 2-h stimulation with TCR Dynabeads, 50 nM ingenol, or 1 µM prostratin. Following stimulation cells were fixed in 4% formaldehyde, permeabilized, and then immunofluorescently co-stained for RasGRP1 and a constitutively expressed membrane protein Ezrin. After counterstaining with DAPI, coverslips were mounted on to glass slides and imaged using a DeltaVision deconvolution microscope at 100X. Scale bar represents a length of 5 µm. *(B)* Degree of membrane colocalization of RasGRP1 and Ezrin as assessed by measuring multiple Pearson’s correlation coefficient values obtained from defined regions of interest. Selected regions of interest defined as single cells were analyzed using the softWoRx colocalization module to measure the degree of colocalization between fluorescently stained RasGRP1 and Ezrin. Statistical significance (*p* values) was calculated using a two-tailed Student’s *t* test. *(C)* UMAP clustering based on scRNA-seq profiling of resting unstimulated and TCR-activated memory CD4^+^ T cells. *(D)* Assessment of the expression profiles of RasGRP isoforms in memory CD4^+^ T cells by scRNA Drop-seq.

In order to determine the expression profiles of each of the four RasGRP isoforms, scRNA- and bulk RNA-seq were carried out in both resting and TCR-activated primary CD4^+^ T cells. **Fig 7C** is a UMAP profiling of memory CD4^+^ T cells that were either unstimulated or activated through the TCR for 24 or 72 h prior to scRNA-seq. By utilizing this algorithmic analytical approach we observed distinct clustering of cells depending on the duration of T-cell activation. Further analysis of these UMAP cell clusters for specific transcript expression revealed that both RasGRP1 and RasGRP2 are expressed at comparable levels in unstimulated memory CD4^+^ T cells with barely any detectable expression of RasGRP3 and RasGRP4 (**Fig 7D**). Irrespective of the duration of TCR co-stimulation, RasGRP3 and RasGRP4 transcripts remained extremely low. Interestingly, RasGRP2 expression was abolished upon TCR activation while RasGRP1 transcript levels were maintained in these cells following 24 h TCR activation but declined slightly at 72 h post-stimulation.

Consistent with these findings, bulk RNA-seq of primary CD4^+^ T cell types that had been polarized into the four major T-cell subsets (Th1, Th2, Th17, and Treg) showed that while RasGRP3 and RasGRP4 mRNA are barely expressed in these cells regardless of the cellular activation state, RasGRP2 transcript expression is highly restricted to cells that are in the quiescent phase (**S19A Fig**). The sharp decline in RasGRP2 mRNA nearly abolished its expression following an overnight period of TCR activation. By contrast, RasGRP1 was found to be expressed at levels that were comparable to those of RasGRP2 in the resting phase of all four CD4^+^ T-cell subsets and despite the greater than 2-fold reduction in RasGRP1 upon overnight TCR co-stimulation, it became the predominantly expressed RasGRP isoform in activated cells (by at least 8-fold higher transcript levels) in view of the near complete abrogation of RasGRP2 and the extremely low expression of RasGRP3 and RasGRP4 (**S19A Fig**).

We performed an additional assessment of RasGRP isoform expression with a bulk RNA-seq dataset generated using memory CD4^+^ T cells that were either unstimulated or subjected to a time course of TCR activation (0, 2, 4, 6, and 24 h) [65]. In agreement with our own scRNA- and bulk RNA-seq experiments described above, analysis of these datasets clearly demonstrated that while RasGRP3 and RasGRP4 transcripts were at very low – at times barely detectable – levels across all time points examined, RasGRP2 expression was comparable to that of RasGRP1 in the unstimulated state then exhibited a rapid and precipitous decay over the 24-h period of T-cell stimulation (**S19B Fig**). By 6 h of TCR activation, RasGRP2 transcript levels had nearly dropped to those of RasGRP3 and RasGRP4. By contrast, RasGRP1 transcript levels sharply declined by greater than 2-fold within 2 h of TCR activation then rose slightly at the 4 h time point before declining again at 24 h (**S19B Fig**). Between the 6 h and 24 h activation period RasGRP1 was clearly found to be the major RasGRP isoform expressed in memory CD4^+^ T cells.

Thus, our RNA-seq analysis reveals that RasGRP1 is by far the predominantly expressed DAG-dependent RasGRP isoform in primary CD4^+^ T cells. Therefore, RasGRP1 is likely to be the isoform that mediates the activation of Ras-Raf-MEK-ERK1/2 signaling in response to TCR co-stimulation or phorbol ester treatment. Furthermore, we have found that mRNA belonging to the DAG-independent isoform RasGRP2 is only present in resting CD4^+^ T cells with its expression being rapidly and nearly abrogated upon T-cell activation.

### PI3K-mTORC1/2 and RasGRP1-ERK1/2 are complementary pathways that are essential for P-TEFb biogenesis and emergence of HIV from latency

Having observed that the MEK inhibitor U0126 causes a modest inhibition of TCR-induced expression of P-TEFb while the Raf inhibitor AZ 628 failed to block the TCR-induced expression of this transcription factor (**Fig 6B and S18 Fig**), we reasoned that there could be other TCR signaling pathways that may contribute to P-TEFb biogenesis in primary T cells. Several years ago, Besnard *et al.* demonstrated that the pharmacological inhibition of mTORC1 and mTORC2 kinase activities blocks the phosphorylation of the CDK9 subunit of P-TEFb in primary CD4^+^ T cells [81]. In the same study it was found that this blockade of CDK9 phosphorylation accounted for the suppression in latent HIV reactivation by the mTOR inhibitors. However, the CDK9 target sites of mTOR-dependent phosphorylation were not defined in the study.

Activation of both mTORC1 and mTORC2 in response to TCR co-stimulation is well established to be PI3K-dependent with AKT being a central mediator of the activating crosstalk between both mTORC complexes (**Fig 8A**) [82, 83]. Since mTORC1 is a critical facilitator of protein synthesis [84], we reasoned that activation of the PI3K-mTORC2-AKT-mTORC1 signaling pathway may contribute to generation of P-TEFb by enabling the translation of the CycT1 subunit which, in line with the P-TEFb biogenesis pathway (**Fig 1A**), is a prerequisite for the subsequent T-loop phosphorylation of CDK9 at both Thr186 and Ser175.

**Fig 8.**
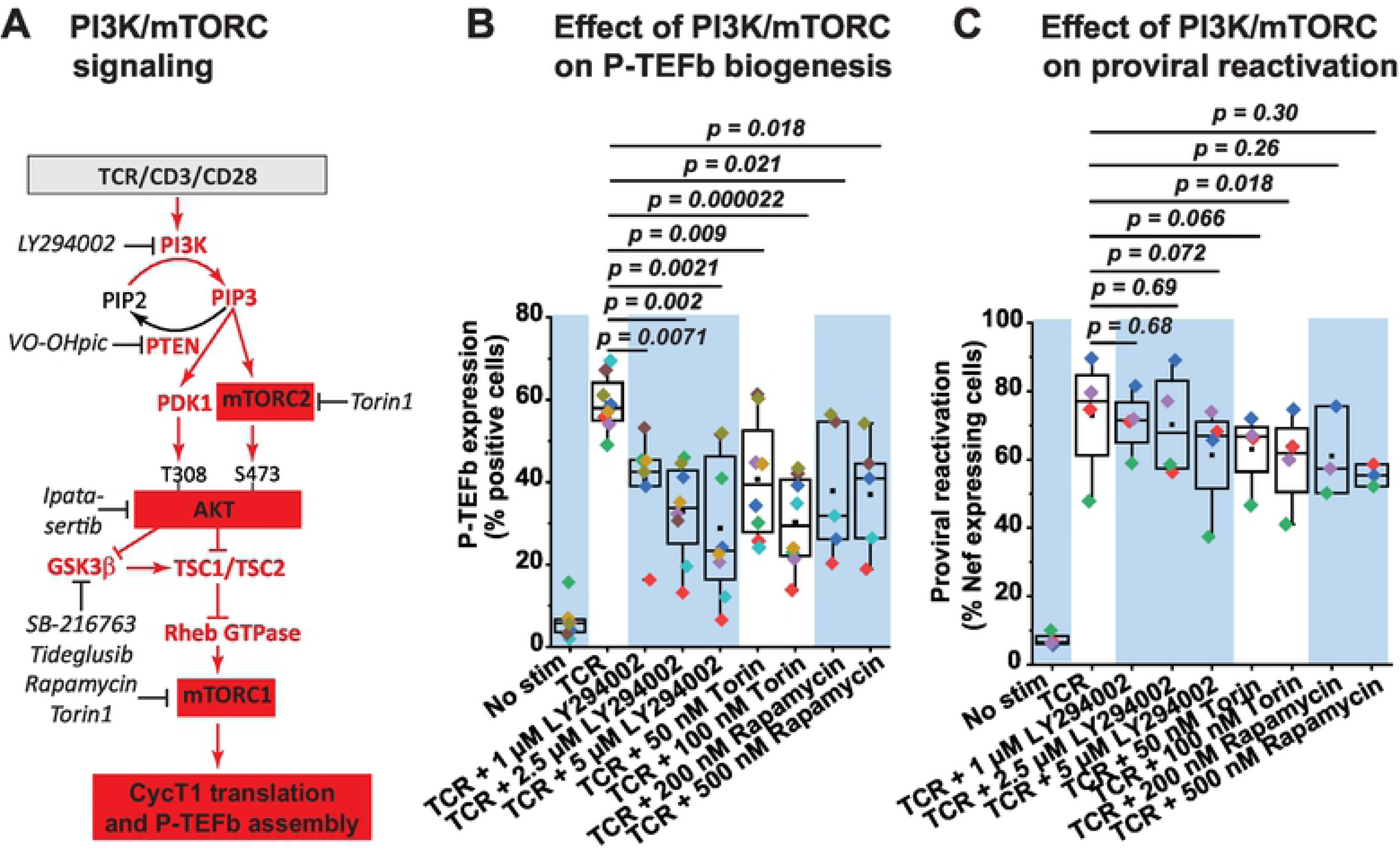
Inhibition of PI3K and mTORC kinases partially suppresses TCR-induced P-TEFb expression and modestly inhibits TCR-induced proviral reactivation in primary CD4^+^ T cells. *(A)* Proposed role of the PI3K/AKT/mTOR signaling pathway in mediating the biogenesis of P-TEFb. Inhibitors that were tested in the current study are shown in black italics. *(B)* PI3K and mTORC kinase inhibitors partially repress TCR-induced expression of active P-TEFb and albeit with significant variability. Memory CD4^+^ T cells (three experiments) and resting primary Th17 cells (four experiments) were treated or not for 30 min with inhibitors targeting PI3K (LY294002), mTORC1 (Rapamycin) or both mTORC1 and mTORC2 (Torin) at the concentrations shown prior to TCR co-stimulation for 24 h. Thereafter, cells were examined for active P-TEFb expression by flow cytometry following dual immunofluorescence staining for cyclinT1 and pSer175 CDK9. *(C)* PI3K and mTORC kinase inhibitors modestly repress proviral reactivation that is in response to TCR co-stimulation. Latently infected Th17 cells prepared using naïve CD4^+^ T cells isolated from three healthy donors were pretreated or not for 30 min with inhibitors to PI3K (LY294002) or mTORC1/mTORC2 (Torin) prior to stimulating the cells through the TCR for 24 h. Cells were thereafter analyzed by flow cytometry following immunostaining using a fluorophore-conjugated antibody towards HIV Nef. Statistical significance (*p* values) in *B* and *C* was determined using a two-tailed Student’s *t* test.

As shown in **Fig 8B**, TCR-induced expression of transcriptionally active P-TEFb was suppressed, albeit with significant variability between experiments, upon treatment of either memory CD4^+^ T cells or polarized Th17 cells with an inhibitor of PI3K (LY294002), mTORC1 (Rapamycin) or a selective mTOR kinase inhibitor (Torin) that blocks both mTORC1 and mTORC2 activities. Mean fold reductions in TCR-induced P-TEFb expression upon PI3K, mTORC1, and mTORC1/2 inhibition were found to be 2.1, 1.9, and 2.2, respectively (**Table 1**). Similarly, inhibition of AKT kinase activity with ipatasertib resulted in a mean fold reduction of 1.7 in TCR-induced P-TEFb expression (**Table 1 and S21A Fig**). Inconsistent with the observed repressive effects on P-TEFb, inhibition of PI3K, mTORC1 and mTORC1/2 kinases elicited very modest reductions in TCR-induced proviral reactivation in primary Th17 cells (**Fig 8C**). Therefore, although PI3K and mTORC inhibitors were found to suppress TCR-induced P-TEFb expression with substantial variability but in a statistically significant manner, they were not as effective at blocking the reactivation of latent HIV.

**Table 1.** Fold reductions in TCR-induced expression of P-TEFb and HIV following inhibition of Raf-MEK-ERK and PI3K-AKT-mTOR pathways.

| ***Kinase(s)*** | ***Inhibitor(s)*** | ***Mean +/- S.E.*** |
| --- | --- | --- |
| **MEK** | 10 μM U0126 | 1.34 +/- 0.17 |
| | 20 μM U0126 | 1.61 +/- 0.22 |
| **Raf** | 100 nM AZ 628 | 0.96 +/- 0.035 |
|  | 500 nM AZ 628 | 1.16 +/- 0.039 |
| **PI3K** | 2.5 μM LY294002 | 2.14 +/- 0.40 |
| | 5 μM LY294002 | 3.19 +/- 0.92 |
| **AKT** | 1 μM ipatasertib | 1.45 +/- 0.17 |
| | 10 μM ipatasertib | 1.72 +/- 0.21 |
| **mTORC1** | 200 nM Rapamycin | 1.90 +/- 0.32 |
|  | 500 nM Rapamycin | 1.93 +/- 0.36 |
| **mTORC1 +**<br>**mTORC2** | 50 nM Torin | 1.62 +/- 0.22 |
|  | 100 nM Torin | 2.20 +/- 0.30 |
| **MEK + PI3K** | 10 μM U0126 + 2.5 μM LY294002 | 5.37 +/- 0.95 |
| | 20 μM U0126 + 2.5 μM LY294002 | 7.83 +/- 3.82 |
| **MEK + mTORC1** | 10 μM U0126 + 200 nM Rapamycin | 5.01 +/- 0.99 |
| | 20 μM U0126 + 200 nM Rapamycin | 8.25 +/- 0.68 |
| **MEK + mTORC1/2** | 10 μM U0126 + 100 nM Torin | 6.57 +/- 1.44 |
| | 20 μM U0126 + 100 nM Torin | 8.73 +/- 1.44 |

Mean values are an average of fold reductions calculated as a ratio of the measure obtained for TCR activation over that obtained for TCR activation in the presence of inhibitor(s). S.E.: standard error of the mean.

Based on these results we reasoned that both the PI3K-AKT-mTORC and the Raf-MEK-ERK pathways may have complementary roles in regulating TCR-induced P-TEFb biogenesis. Consistent with this idea, we found that a combined inhibition of MEK and PI3K or MEK and mTORC in both primary Th17 and memory CD4^+^ T cells was much more effective at preventing P-TEFb expression upon TCR co-stimulation than inhibition of the individual kinases (**Figs 9A, 9B and S22 Fig, Table 1**). Therefore, the combined treatment of these cells with U0126 and LY294002 or U0126 and either mTORC inhibitor (rapamycin for mTORC1 or Torin for mTORC1 and mTORC2) nearly abrogated TCR-induced P-TEFb expression and with minimal cytotoxicity (**Figs 9A and 9B**).

**Fig 9.**
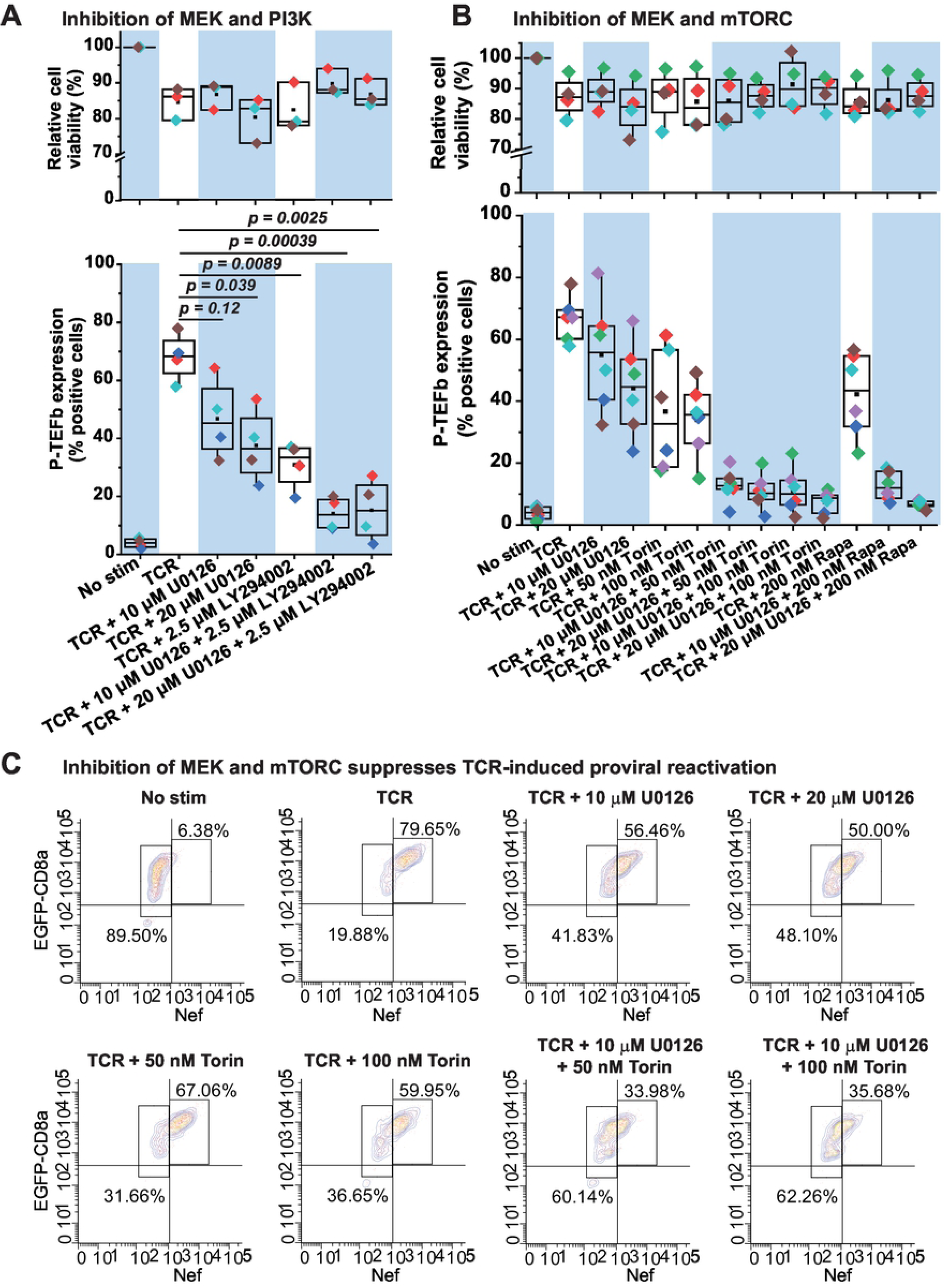
Combined inhibition of MEK and mTORC kinases abrogates TCR-induced P-TEFb expression and effectively suppresses TCR-mediated proviral reactivation in primary CD4^+^ T cells. *(A)* Combined inhibition of MEK and PI3K kinases enhances the disruption of the active P-TEFb expression. Healthy donor memory CD4^+^ T cells (in two of the plotted experiments) and polarized resting primary Th17 cells (in two of the plotted experiments) were treated or not for 30 min with inhibitors targeting either MEK (U0126) or PI3K (LY294002) or both at the concentrations shown prior to TCR co-stimulation for 24 h. Thereafter, cells were examined for active P-TEFb expression by flow cytometry following dual immunofluorescence staining for cyclinT1 and pSer175 CDK9. Cell viability was also assessed by flow cytometry in three of the four experiments following staining with the eFluor450 viability dye. The viability data shown are expressed as a percentage relative to non-treated cells. *(B)* Combined inhibition of MEK and mTORC abrogates the expression of active P-TEFb. Memory CD4^+^ T cells (four experiments) and resting primary Th17 cells (two experiments) were treated or not for 30 min with the MEK inhibitor U0126 on its own or in combination with inhibitors targeting mTORC1 (Rapamycin) or both mTORC1 and mTORC2 (Torin) at the concentrations shown prior to TCR co-stimulation for 24 h. Thereafter, cells were examined for active P-TEFb expression by flow cytometry following dual immunofluorescence staining for cyclinT1 and pSer175 CDK9. In four of the experiments shown, cell viability was assessed by flow cytometry following staining with the eFluor450 viability dye. The viability data are expressed as a percentage relative to non-treated cells. Statistical significance (*p* values) in *A* and *B* was determined using a two-tailed Student’s *t* test. *(C)* Latently infected Th17 cells were pretreated or not for 30 min with the MEK inhibitor U0126 or the mTORC1/2 inhibitor Torin on their own or in combination prior to TCR co-stimulation for 24 h. Afterwards, cells were analyzed by flow cytometry following immunostaining using a fluorophore-conjugated antibody towards HIV Nef.

We also examined the effect of these combined treatments on proviral expression following TCR co-stimulation of latently infected primary Th17 cells. Inhibition of both mTORC1 and mTORC2 with Torin was most effective in combination with U0126 at suppressing proviral reactivation beyond that observed with either inhibitor (**Fig 9C and S23 Fig**). Overall, these results indicate that two of the major signaling arms of the TCR cascade, namely, RasGRP1–Ras–Raf– MEK–ERK1/2 and PI3K–mTORC2-AKT–mTORC1 serve essential complementary functions in mediating the biogenesis of P-TEFb in primary CD4^+^ T cells that is in turn crucial to enabling the emergence of HIV from latency.

## Discussion

### Regulation of P-TEFb in primary T cells is central to HIV transcriptional control

Dissecting the cellular pathways through which HIV emerges from latency is critical to the development of “shock and kill” therapeutic approaches that can be used to effectively eradicate persistent reservoirs of transcriptionally latent but replication-competent HIV proviruses in memory CD4^+^ T cells of infected individuals. Using the well-characterized and highly reproducible primary QUECEL model of HIV latency together with healthy donor memory CD4^+^ T cells, we identified a series of T-cell signaling pathways that are crucial to the emergence of HIV from latency. In particular, the host transcription elongation factor P-TEFb is an obligate activator of proviral HIV transcription, and therefore, comprehensively understanding its regulation by signal transduction mechanisms in primary T cells is central to HIV cure efforts.

P-TEFb is expressed at vanishingly low levels in memory CD4^+^ T cells and this is one of the key regulatory mechanisms that minimizes HIV transcription in resting cells. However, TCR co-stimulation initiates a set of signaling cascades that result in the posttranscriptional expression of its CycT1 subunit and T-loop phosphorylation of CDK9 kinase at Ser175 and Thr186. Therefore, the studies presented here were specifically aimed at identifying precise TCR signaling pathways in primary T cells that are essential for the generation of transcriptionally active P-TEFb, defined as the coordinate expression of CycT1 and pSer175 CDK9.

Our major findings, as summarized in the scheme shown in **Fig 10**, are: i) reactivation of HIV from latency is tightly coupled to the biogenesis of P-TEFb and both of these processes can be elicited upon cellular treatment with PKC agonists in the absence of intracellular calcium mobilization with an ionophore stimulus; ii) stimulation of both P-TEFb expression and proviral reactivation upon TCR co-stimulation or challenge with PKC agonists is largely independent of PKC activity; iii) PKC agonists signal through off-target activation of the RasGRP1-Ras-Raf-MEK-ERK1/2 to stimulate both P-TEFb biogenesis and the reactivation of latent HIV; and iv) generation of transcriptionally active P-TEFb upon TCR co-stimulation is primarily mediated by two complementary pathways, namely, the RasGRP1-Ras-Raf-MEK-ERK1/2 and the PI3K– mTORC2-AKT–mTORC1 pathways.

**Fig 10.**
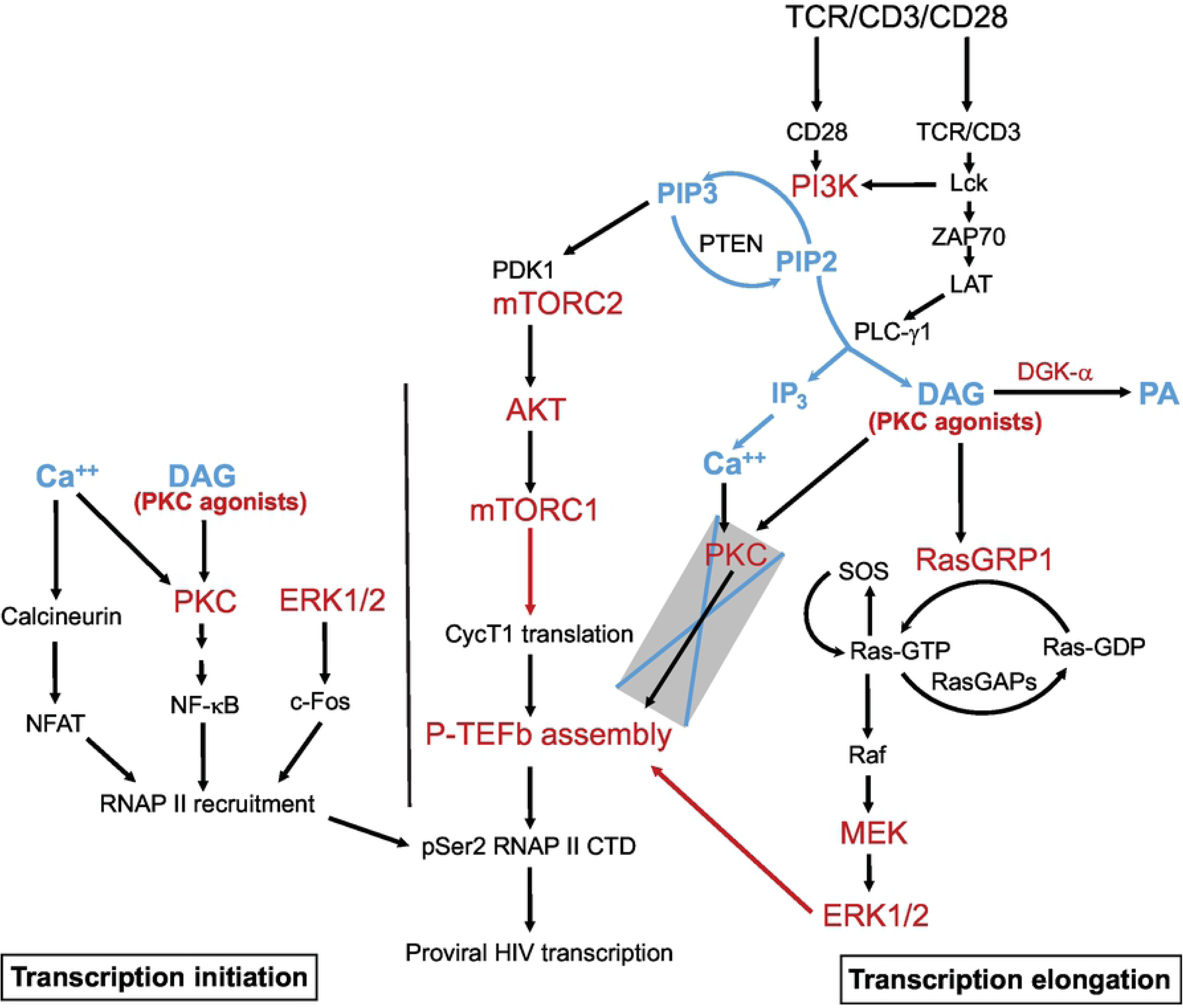
Model for the reactivation of P-TEFb and proviral HIV by RasGRP1-Ras-Raf-MEK-ERK1/2 and PI3K-AKT-mTORC1/2 signaling in latently infected CD4^+^ T cells. Generation of diacylglycerol (DAG) by activated PLC-γ1 immediately following TCR co-stimulation enables the recruitment of RasGRP1 to the plasma membrane, the initial activation of Ras via GTP loading and the allosteric activation of membrane-anchored SOS by GTP-bound Ras. This can also be accomplished by DAG-mimicking PKC agonists. Allosteric activation of SOS creates a positive Ras-GTP-SOS feedback loop that leads to maximal activation of Ras and formation of Ras-Raf complexes at the membrane. A phosphorylation cascade is triggered by Ras-Raf leading to activation of ERK1 and ERK2 which is turn stimulate the biogenesis of P-TEFb via mechanisms that are yet to be defined. RasGRP1 activity may be regulated by controlling the availability of DAG which can get converted to phosphatidic acid (PA) primarily by DGK-α. TCR co-stimulation also rapidly stimulates PI3K leading to the activation of mTORC1 which may regulate P-TEFb biogenesis by enabling posttranscriptional synthesis of cyclin T1 (CycT1). Inhibitor-based experiments have clarified that intracellular calcium release and the activation of PKC are dispensable for formation of transcriptionally active P-TEFb. However, the recruitment of RNA polymerase II (RNAP II) to proviral HIV and its eventual phosphorylation by P-TEFb to stimulate processive transcription elongation are likely to be dependent on both Ca^++^ and PKC signaling.

### Off-target pathway mediating P-TEFb induction by PKC agonists

T-cell challenge with a PKC agonist (usually PMA) plus a calcium ionophore has been widely used in the HIV cure field as a gold standard for measuring the relative effectiveness of LRAs at reversing HIV latency in both *in vitro* primary cell models and patient-derived CD4^+^ T cells [85–87]. In our current study using the QUECEL primary T-cell HIV latency model, we found that single challenge with ingenol or prostratin reproducibly elicited levels of proviral reactivation that were comparable to TCR co-stimulation. Each of the PKC agonists we examined was able to stimulate active P-TEFb expression in both primary Th17 and memory T cells, albeit not as effectively as TCR co-stimulation. However, co-treatment with ionomycin failed to improve their stimulatory effects on P-TEFb. Moreover, ionomycin treatment on its own at best modestly induced CycT1 and pSer175 CDK9 expression. These findings strongly suggest that the calcium signaling arm of the TCR pathway may not crucial for the inducible expression of P-TEFb in these cells.

Calcium signaling has been shown to be essential for the initiation of HIV transcription in the QUECEL model via activation of NFAT [64]. However, it seems likely that under conditions of phorbol ester treatment alone HIV transcription initiation is mediated primarily by NF-κB as is seen in the Jurkat T-cell models [35, 88, 89]. Co-treatment of memory and Th17 cells with ionomycin significantly potentiated the efficacy of PKC agonists at generating the transcriptionally active form of NF-κB (pSer529 p65) and the transcription elongation competent form of RNAP II (pSer2 CTD RNAP II), which provides a downstream readout of P-TEFb enzymatic activity, since pSer2 CTD RNAP II is a modification that is at least in part due to P-TEFb and is strictly associated with polymerase that is already present on transcribing genes [90, 91]. It is therefore quite likely that calcium signaling serves to promote the recruitment of RNAP II to gene promoters in manner that is dependent upon the activation of both NF-κB and NFAT. It is also noteworthy that single challenges with the PKC agonists were minimally cytotoxic to primary T cells while the co-treatment with ionomycin resulted in a significant loss of cell viability. These observations are consistent with a recent report showing that treatment of CD4^+^ T cells with prostratin or byrostatin-1 induces resistance to apoptosis in a PKC-dependent manner [92]. Thus, because of their ability to induce P-TEFb biogenesis and stimulate the reactivation of HIV from latency, more selective PKC agonists show strong potential as candidate LRAs.

It is generally believed that the latency reversal activity of PKC agonists, also commonly referred to as phorbol esters is due to the activation of PKC-dependent pathways. These agents structurally mimic endogenously generated DAG to bind the C1 domains of both conventional (α, βI, βII, and γ) and novel (δ, θ, ε, and η) PKC isozymes leading to their membrane recruitment and stimulation of their kinase activities [30]. However, they can also target several other proteins with C1 domains including RasGRP, a critical guanine nucleotide exchange factor for Ras in T cells, α and β chimaerins, Munc13 proteins, protein kinase D (PKD) and DAG kinases (DGKs) [93]. In our current study, we unexpectedly found that pan-PKC inhibitors (which potently inhibit α, β, γ, δ, and ε isoforms) and an inhibitor that selectively targets PKC-θ, considered to be the predominantly functional PKC isoform in T cells [68–70], were all ineffective at blocking phorbol ester- or TCR-induced P-TEFb expression and proviral reactivation in primary T cells. By contrast, selective inhibitors of Raf and MEK kinases effectively suppressed the induction of P-TEFb that was in response to phorbol ester treatment. Moreover, the MEK inhibitor also effectively suppressed phorbol ester-induced proviral reactivation in the QUECEL model. These results strongly indicate that these agents are signaling through the Ras-Raf-MEK-ERK pathway rather than through PKC in eliciting their stimulatory activity on P-TEFb and HIV in primary T cells.

### RasGRP1 is a key mediator of P-TEFb biogenesis

A critical initiator of Ras-Raf-MEK-ERK MAPK signaling in response to cellular treatment with DAG-mimicking phorbol esters is the Ras GDP-GTP exchange factor RasGRP [29, 74]. Of the four known isoforms of RasGRP, RasGRP1, RasGRP3 and RasGRP4 have been shown to bind phorbol esters with high affinity via their C1 domains and, consequently, they get recruited to the plasma membrane where they are able to access Ras and facilitate guanine nucleotide exchange [29, 75–77]. SOS, another important Ras GDP-GTP exchange factor, does not bind DAG but is rather anchored to the membrane by a complex of tyrosine-phosphorylated LAT and the adaptor protein Grb2 [94]. Following T-cell stimulation guanine nucleotide exchange activity of membrane-associated SOS is triggered by the initial activation of Ras by RasGRP [71]. In turn GTP-bound Ras allosterically binds SOS leading to its full activation. This creates a positive Ras-GTP-SOS feedback loop that results in a conformational change in Ras which permits robust formation of Ras-Raf complexes at the membrane [71, 95].

Transcriptome analyses of QUECEL T-cell subsets and memory CD4^+^ T cells performed in our study using scRNA- and bulk RNA-seq approaches, unequivocally demonstrated that of the three DAG-dependent RasGRP isoforms, RasGRP1 is by far the predominantly expressed isoform in resting and TCR-activated cells. By contrast, RasGRP3 and RasGRP4 transcripts were barely detected under both conditions. Assuming mRNA copy numbers of these isoforms appropriately reflect their relative protein expression and potential activities, it is quite likely that the activation of ERK MAPK signaling following TCR co-stimulation or challenge with PKC agonists is being mediated by RasGRP1. Furthermore, we were able to confirm by immunofluorescence microscopy that RasGRP1 rapidly translocates to the membrane following stimulation of memory T cells with PKC agonists. Therefore, these overall results implicate RasGRP1 as being the Ras GDP-GTP exchange factor that orchestrates the regulation of P-TEFb biogenesis in primary T cells.

Unlike the DAG-dependent RasGRP proteins, RasGRP2 has a very poor binding affinity for DAG and fails to translocate to the plasma membrane in response to treatment with DAG analogs [78–80]. Membrane trafficking of RasGRP2 has been shown to be dependent on F actin polymerization [96]. Moreover, a longer splice variant of RasGRP2 is found to be myristoylated and palmitoylated at its N-terminus which also enables a DAG-independent membrane localization [97]. Our transcriptome studies showed that RasGRP2 transcripts are highly restricted to resting primary T cells. Although RasGRP2 mRNA expression was initially comparable to that of RasGRP1 in the unstimulated state, it rapidly and precipitously decayed following TCR activation eventually reaching the extremely low levels seen with RasGRP3 and RasGRP4. Dramatic changes in the abundance of a transcript tend to correlate strongly with changes in the abundance of its protein as we have observed in the case of ELL2 and HEXIM1 (**S6 Fig**). Therefore, these RNA-seq results could be an indication that RasGRP2 is functionally relevant in the context of resting T cells or during the earlier period of T-cell activation.

Our findings are noteworthy in view of the established role of RasGRP2 as the primary activator of the Ras homolog Rap1 [97, 98], which has been demonstrated in multiple studies to be capable of antagonizing Raf-activating Ras activity in T cells [99–101]. Interestingly, TCR stimulation of primary human T cells with anti-CD3 in the absence of co-stimulatory signals was shown to rapidly and transiently activate Rap1 (within 10 min of receptor crosslinking with anti-CD3) while co-stimulation with anti-CD28 abolished the anti-CD3-dependent activation of Rap1 [102]. As shown in **S19C and S19E Figs**, both unstimulated memory T cells and QUECEL T-cell subsets expressed much higher copy numbers of Rap1A and Rap1B transcripts as compared to those belonging to H-Ras, K-Ras and N-Ras. These observations warrant further research because they hint at a possible RasGRP2-Rap1 functional interaction that may modulate formation of P-TEFb perhaps by counteracting RasGRP1-Ras-Raf signaling in primary T cells.

### Combined roles of the RasGRP1-Ras-Raf-MEK-ERK1/2 and PI3K-mTORC2-AKT-mTORC1 pathways in mediating P-TEFb biogenesis

Molecular mechanisms through which the RasGRP1-Ras-Raf-MEK-ERK1/2 pathway effects the formation of transcriptionally active P-TEFb in primary T cells are still yet to be defined. Using a Jurkat T-cell model system for HIV latency, we had previously shown that ERK MAPK signaling in response to phorbol ester or TCR co-stimulation can serve as a primary mediator of the dissociation of P-TEFb from 7SK snRNP [36]. By conducting chromatin immunoprecipitation studies we were able to further demonstrate that inhibition of ERK activation by MEK substantially suppressed HIV transcription elongation by preventing the recruitment of P-TEFb to latent proviruses. MEK inhibition also suppressed the T-loop phosphorylation of CDK9 at Ser175 in Jurkat T cells, a modification that is exclusively found on P-TEFb that is not associated with 7SK snRNP [60]. In both of these studies we were unable to investigate whether this pathway is also essential for the biogenesis of P-TEFb since Jurkat T cells constitutively express steady-state levels of the enzyme, the majority of which is already assembled into 7SK snRNP.

Since the posttranscriptional synthesis of CycT1 is essential for initiating the assembly of enzymatically active P-TEFb, as proposed in **Fig 1**, determining the exact mechanisms by which ERK MAPK signaling orchestrates the translation of CycT1 in primary T cells will be of paramount importance in our future studies. Two possible mechanisms may involve relieving the microRNA-directed restriction of CycT1 protein synthesis in resting T cells or facilitating the nuclear export of the RNA binding protein NF90. The latter was previously shown to be capable of binding the 3’ untranslated region of CycT1 mRNA in latently infected monocyte-derived U1 cells in order to promote translation initiation of CycT1 and subsequent induction of proviral HIV [103].

Our identification of PI3K–mTORC2-AKT–mTORC1 as a complementary TCR pathway for stimulating P-TEFb expression suggests that mTORC1 signaling can participate in P-TEFb biogenesis by enhancing the translation of CycT1 mRNA. In mammalian cells, stimulation of mTORC1 activity is critical to the rapid initiation of protein synthesis through the sequential phosphorylation and activation of components of the translational machinery which include the ribosomal protein S6 kinase 1 (S6K1), its downstream substrate ribosomal protein S6 as well as initiation and elongation factors [104]. Importantly, there is a precedent for a crosstalk between ERK and mTORC signaling leading up to the activation of these translation factors. A study investigating the basis for cardiac hypertrophy reported that Gq protein-coupled receptor agonists induce protein synthesis in cardiomyocytes by stimulating the phosphorylation of S6K1 and eIF4E-binding protein-1 (4E-BP1) via Ras-ERK signaling rather than through AKT and this ability is effectively blocked either by inhibition of MEK or mTORC1 [105]. It will therefore be interesting to examine whether stimulation of ERK activity in primary T cells with PKC agonists may in turn lead to stimulation of mTORC1 and phosphorylation of translation factors.

Disulfiram (DSF), an inhibitor of acetaldehyde dehydrogenase that is used for the treatment of alcoholism, was shown to reactivate latent HIV-1 expression in a primary cell model of virus latency and functions by depleting PTEN protein expression [106]. PTEN lipid phosphatase and GSK3β kinase are well established negative regulators of the PI3K–mTORC2-AKT–mTORC1 signaling pathway (**Fig 8A**) [107, 108]. However, in our study we found that inhibition of their activities either individually or in tandem was not sufficient to elicit P-TEFb expression in memory T cells (**S21 Fig**). Therefore, in future studies we will be testing to see whether various combination treatments that involve using inhibitors of PTEN, GSK3β, diacylglycerol kinase-α (DGK-α; which as shown in **S20A** and **S20B Figs** is the predominant DGK isoform in primary CD4^+^ T cells) as well as membrane-permeable diacylglycerols can stimulate P-TEFb biogenesis.

## Conclusions

Despite extensive small molecule screens, it has been challenging to identify potent latency reversing agents. It seems likely that this is because there are multiple barriers to HIV reactivation in latently infected cells involving both transcription initiation, transcription elongation and epigenetic regulation. Despite this complexity, the major rate-limiting step for HIV transcription appears to involve P-TEFb activation. Our studies have shown that although P-TEFb biogenesis after TCR stimulation is the result of multiple complementary signaling pathways, effective activation can be achieved through the RasGRP1-Raf-ERK1/2 pathway. It therefore seems likely that selectively activating this pathway in combination with activation of a transcription initiation factor, for example using a non-canonical activator of NF-κB [109], provides an optimal strategy for latency reversal in the absence of full T-cell activation – the key missing element in the “shock and kill” HIV eradication strategy.

## Materials and Methods

### Materials

RPMI 1640 medium and fetal bovine serum for culturing memory CD4^+^ T cells and preparing polarized Th17 cells were purchased from Hyclone. Human T-cell activator anti-CD3 and anti-CD28 antibody cocktail covalently bound to magnetic beads (Dynabeads) was obtained from ThermoFisher Scientific. Soluble anti-CD3 and anti-CD28 antibodies were from BD Biosciences. Ingenol-3-angelate and SAHA were purchased from Cayman Chemical. Recombinant TNF-α was from R&D Systems. Prostratin, PMA and ionomycin were from Millipore Sigma. The sources and catalog information of all commercially obtained inhibitors and antibodies used in the current study are shown in **S1 Table**. HEXIM1 antibody was custom-synthesized and affinity purified by Covance Research Products. Generation, purification and validation of the polyclonal antibody towards the phospho-Ser-175 CDK9 epitope has been described previously [60].

### Isolation and experimental culture of memory CD4^+^ T cells

Approximately 1.0 × 10^7^ memory CD4^+^ T cells were routinely isolated by negative bead selection from previously frozen healthy donor leukapheresis packs containing 1.0 × 10^8^ peripheral blood mononuclear cells (PBMCs) using the EasySep™ memory CD4^+^ T-cell enrichment kit (StemCell). After isolation cells were allowed to recover overnight in complete RPMI media lacking IL-2 prior to treating them with inhibitors and/or stimuli described in the current study. Purity and resting status of the isolated memory T cells were confirmed by immunofluorescence flow cytometry analysis of CD45-RO, cyclin D3 and Ki67 expression. For the inhibitor experiments, memory T cells were equally divided into a 24-well plate at approximately 3.0 × 10^5^/ml of complete media and pretreated with the appropriate inhibitors for 30 min before challenging them with either anti-CD3/anti-CD28 Dynabeads at a 1:1 bead-to-cell ratio or with the phorbol esters ingenol-3-angelate, prostratin or PMA. For cell viability assessment, experimentally treated cells were harvested by centrifugation at 1500 rpm for 3 min in polystyrene 5-ml round-bottomed flow cytometry tubes, washed with 1X PBS and then subjected to staining with eFluor450 Viability Dye (ThermoFisher Scientific) for 30 min on ice.

### Preparation of quiescent and HIV-infected Th17 cells

The procedure for generating quiescent and/or latently infected primary Th17 cells has been extensively described previously [64]. Briefly, naïve CD4^+^ T cells isolated from previously frozen healthy donor leukapheresis packs by negative magnetic bead selection using the EasySep™ Naïve CD4^+^ T-cell isolation kit (StemCell) were treated for 8 days with 10 μg/ml Concavanalin A and a cocktail of polarizing cytokines and antibodies. On Day 4, IL-2 was added to the cells at 60 IU/ml. Following Th17 polarization and T-cell expansion, cells were infected with a VSVG-pseudotyped pHR’-Nef+-CD8a/GFP viral construct by spinoculation. On Day 14, HIV infected cells were isolated by anti-CD8a magnetic separation. To generate quiescent T cells, both the infected and the uninfected (negatively sorted) cells were cultured in medium containing reduced concentrations of IL-2 (15 IU/ml) and IL-23 (12.5 µg/ml) for at least 2 weeks. Entry into quiescence was monitored by flow cytometry by measuring EdU incorporation and analyzing the expression of cyclin D3 and Ki67. Proviral latency was also monitored by assessing the expression of Nef by immunofluorescence flow cytometry before and after TCR reactivation for 24 h.

### Immunofluorescence flow cytometry

Experimentally treated primary Th17 or memory CD4^+^ T cells were harvested by centrifugation at 1500 rpm for 3 min in polystyrene 5-ml round-bottomed flow cytometry tubes. Thereafter, cells were washed with 1X PBS prior to fixation in 4% formaldehyde for 15 min at room temperature. For surface-staining with fluorophore-conjugated CD45-RO, CCR7, CD25 and CD69 antibodies, fixed cells were first washed with 1X PBS then incubated with these antibodies for 45 min in the dark and at room temperature. For intracellular staining, fixed cells were permeabilized by a sequential treatment with 0.2% Triton-X-100 for 10 min and incubation with 1X BD Perm/Wash buffer (BD Biosciences) for 15 min at room temperature. Following a 15-min blocking step with a non-specific IgG, permeabilized cells were immunostained for 45 min in the dark and at room temperature with fluorophore-conjugated antibodies against cyclin D3, Ki67, total CDK9, cyclin T1, pThr186 CDK9, pSer175 CDK9, pSer2 CTD RNA polymerase II, or pSer529 p65 NF-κB. For antibodies that were unconjugated, Alexa Fluor™ antibody labeling kits from ThermoFisher Scientific were used for fluorophore conjugation. After immunostaining cells were rinsed with 1X PBS and then subjected to flow cytometry analysis using the BD LSR Fortessa instrument (BD Biosciences) equipped with the appropriate laser and filters.

### Immunofluorescence deconvolution microscopy

Approximately 3.0 × 10^5^ experimentally treated memory CD4^+^ T cells were harvested by centrifugation at 1500 rpm for 3 min, washed with 1X PBS and resuspended in 30 µl 1X PBS. Resuspended cells were then placed onto poly-L-lysine-coated coverslips in a 24-well plate and allowed to adhere to the surface at 37°C for 15 min. Thereafter, cells were fixed with 4% formaldehyde for 15 min, washed with 1X PBS and then permeabilized with a sequential treatment of 0.2% Triton-X-100 for 10 min followed by 1X Perm/Wash buffer for 30 min at room temerature. Blocking was performed using 10% Normal Donkey Serum (The Jackson Laboratory) prepared in 1X BD Perm/Wash for 30 min and, afterwards, cells were immunostained for 2 h at room temperature with primary antibodies against CDK9, Cdc37, cyclin T1, RasGRP1 or Ezrin. Primary antibody solutions were all prepared in blocking buffer. Thereafter, cells were washed three times with 1X Perm/Wash buffer and then incubated for 1 h with 1:200 dilutions of fluorophore conjugated anti-rabbit or anti-mouse IgG secondary antibodies in blocking buffer. After washing the coverslips with 1X Perm/Wash buffer, they were counterstained with 1 μg/ml DAPI for 15 min, mounted onto glass slides with ProLong Antifade medium (ThermoFisher Scientific), and imaged at high magnification (100X) using a DeltaVision epifluorescent microscope (Applied Precision). Images were captured in a z stack, deconvolved and processed using the softWoRx analysis program (Applied Precision). For RasGRP1 and Ezrin immunostaining, the softWoRx colocalization module was used to create scatterplots and measure the Pearson’s correlation coefficient. A region of interest for the colocalization analysis was defined as a single cell. Processed images were exported as JPEG files and micrographs were composed using Adobe Photoshop.

### Combined HEXIM1 immunofluorescence and 7SK RNA FISH staining

Memory CD4^+^ T cells that were washed and resuspended in 1X PBS as described above were placed onto poly-L-lysine-coated coverslips in a 24-well plate and allowed to adhere to the surface at 37°C for 15 min. Thereafter, cells were fixed with 4% formaldehyde for 15 min, washed with 1X PBS and then permeabilized with a sequential treatment of 0.2% Triton-X-100 for 10 min followed by 1X Perm/Wash buffer for 30 min. For the blocking step, cells were incubated in blocking solution for 30 min that was prepared as follows: 1% BSA (10 µg/ml) and 50 µg/ml Donkey IgG in 1X Perm/Wash. This blocking buffer was also used to prepare the primary and secondary antibody solutions. After immunostaining the cover slips were washed twice with 1X PBS and then fixed in 4% formaldehyde solution for 15 min at room temperature. Coverslips were washed once with 1X PBS prior to incubation with Wash Buffer A (2X SSC and 10% formamide in RNase-free ultrapure H_2_O) for 5 min at room temperature. For RNA FISH, fluorophore-conjugated oligonucleotide probe was resuspended in hybridization buffer (10% dextran sulfate, 2X SSC and 10% formamide in RNase-free ultrapure H_2_O) to a final concentration of 125 nM. Coverslips were inverted onto 40 μl droplets of the hybridization mixture in a humidified chamber which was then tightly sealed and incubated in the dark at 37°C for 16 hours. Thereafter, coverslips were transferred onto a fresh 24-well plate cells side up, incubated in the dark with wash buffer A for 30 min and then counterstained for 15 min with 1 μg/ml DAPI that was prepared in Wash Buffer A. Afterwards, the coverslips were washed twice with Wash Buffer B (2X SSC in RNAase-free ultrapure H_2_O) prior to mounting onto glass slides using Prolong Antifade medium. Images were captured at high magnification (100X) by deconvolution microscopy and then processed as described above.

### Subcellular fractionation of primary Th17 cells for Western blotting

Low-salt and chromatin nuclear fractions were prepared as described previously [25]. Briefly, unstimulated and TCR-activated primary Th17 cells were harvested and washed with ice-cold 1X PBS. Cells were then resuspended in low-salt Cytosolic Extract (CE) lysis buffer (10 mM KCl, 10 mM MgCl2, 0.5% NP-40, 1 mM DTT, and 10 mM Hepes, pH 7.8) containing an EDTA-free protease and phosphatase inhibitor mixture (Roche Applied Science) and placed on ice for 30 min. Thereafter, nuclei were gently pelleted by centrifugation at 4000 rpm for 5 min and washed once with CE buffer. The nuclear pellets were placed in Cell Lysis Buffer A (150 mM NaCl, 10 mM KCl, 1.5 mM MgCl_2_, 0.5% NP-40, 1 mM DTT, 10 mM Hepes pH 8.0) containing a cocktail of protease and phosphatase inhibitors and lysed in multiple quick freeze/thaw cycles using isopropyl alcohol/dry ice and a 37°C water bath. This was followed by sonication using a microtip to mechanically shear DNA and solubilize chromatin-bound proteins. Nuclear extracts were then centrifuged at 4000 rpm for 3 min to remove insoluble debris. Protein concentration in both low-salt and chromatin nuclear fractions was determined at 280 nm using a Nanodrop spectrophotometer (ThermoFisher Scientific). Thereafter, both fractions were subjected to Western blotting using primary antibodies towards AFF1, ELL2, ENL, CDK9, cyclin T1, HEXIM1, p65, p50, NFAT-c1, c-Fos and c-Jun.

### Single-cell transcriptome profiling of memory CD4^+^ T cells

Memory CD4^+^ T cells were isolated from a healthy donor leukapheresis pack and allowed to recover overnight in complete RPMI medium with no IL-2 addition prior to challenging them or not with anti-CD3/anti-CD28 Dynabeads for 24 or 72 h at a 1:1 bead-to-cell ratio in media containing 15 IU/ml IL-2. Thereafter, cells were prepared for Drop-seq by magnetically removing the T-cell receptor activator beads, removal of dead cells using the MACS dead cell removal kit and resuspension of cells in 1X PBS containing 0.1% BSA at 2 × 10^5^ cells/ml. To perform transcriptome profiling, a total of 600,000 single cells from each of the three samples were subjected to Drop-seq. Briefly, cells were resuspended in PBS/1% BSA at 200,000 cells/ml. Drop-seq was performed as previously described [110] using microfluid device (FlowJEM, Toronto, Canada) and barcoded beads (ChemGenes, Inc.). After the isolation of single cells with barcoded beads in oil droplets (Bio-Rad, Inc.), we prepared indexed cDNA libraries using Nextera XT library preparation kit (Illumina, Inc.). Libraries were sequenced using Illumina HiSeqX (Medgenome, Inc.). We mapped the resulting reads against the human GRCh38.p13 reference genome and used the Drop-Seq Toolkit v. 1.0 to generate the digital gene expression (DGE) matrices. Our primary data is publicly available at GEO (Accession no. GSE167916). We analyzed, visualized and explored the resulting DGE matrices using an open-source Seurat software package for R, version 4 [111–113]. Each DGE matrix was used to generate a Seurat object. Each single-cell transcription profile taken for further analysis expressed at least 100 genes. We used the SCTransform protocol to normalize the data. Common anchor features (genes) were identified and used to combine the data from different samples into a single Seurat object. After the identification of highly variable genes, principal component analysis (PCA) was performed. Significant principal components were determined using the Jackstraw plot analysis. To classify distinct groups of cells we performed the dimensionality reduction using the Uniform Manifold Approximation and Projection (UMAP) algorithm [114]. Cells were clustered according to their unbiased transcriptome signatures (unsupervised clustering). We used the Dot-plot function in Seurat to quantify positive cell enrichment and relative expression levels of genes of interest in the dataset.

### Bulk RNA-seq of primary CD4^+^ T-cell QUECEL subsets

Bulk RNA-seq analysis was performed on cells in fully quiescent state and 24 hours following reactivation according to the QUECEL protocol [64] (SRA accession SRP145508). RNA-seq reads were quality controlled using Fastqc and trimmed for any leftover adaptor-derived sequences, and sequences with Phred score less than 30 with Trim Galore, which is a wrapper based on Cutadapt and FastQC. Any reads shorter than 40 nucleotides after the trimming was not used in alignment. The pre-processed reads were aligned to the human genome (hg38/GRCh38) with the Gencode release 27 as the reference annotations using STAR version 2.7.2b [115], followed by gene-level quantitation using htseq-count [116]. In parallel, the pre-processed reads were pseudoaligned using Kallisto version 0.43.1 [117] with 100 rounds of bootstrapping to the Gencode release 27 of the human transcriptome to which the sequence of the transfected HIV genome and the deduced HIV spliced transcripts were added. The resulting quantitations were normalized using Sleuth. The two pipelines yielded concordant results. Transcripts per million (TPM) values were used to evaluate the relative abundance of transcripts under resting and activated conditions. A publicly available bulk RNA-seq dataset of primary human memory CD4^+^ T cells that had been activated or not through TCR co-stimulation (SRA accession SRP026389) was subjected to an identical analysis workflow as described above.

## Acknowledgements

We thank all past and present members of the Karn laboratory for their help and useful discussions. We also thank the CWRU/UH Center for AIDS Research for the provision of microscopy and flow cytometry services.

## Supporting information

**S1 Fig. Isolation of memory CD4^+^ T cells and their characterization.** *(A)* Surface phenotype of memory CD4^+^ T cells isolated using the EasySep™ Memory CD4^+^ T Cell Enrichment Kit (Cat. # 19165). Following memory CD4^+^ T-cell isolation cells were treated or not for 24 h with the stimuli shown then subjected to immunofluorescence flow cytometry after staining with fluorophore-conjugated isotype controls or antibodies towards CD25 and CD45-RO. (B) Central memory CD4^+^ T cells account for the majority of both total peripheral CD4^+^ and memory CD4^+^ T cells. Total peripheral CD4+ T cells were isolated from healthy donor PBMCs using EasySep™ Human CD4^+^ T Cell Kit (Cat. # 17952). Thereafter, cells were subjected to immunofluorescence flow cytometry after staining with fluorophore-conjugated antibodies towards CCR7 and CD45-RO. Based on this dual staining, CD4^+^ T-cell subsets were determined in multiple experiments as a fraction of total peripheral CD4^+^ or total memory CD4^+^ T cells.

**S2 Fig. Early expression of P-TEFb coincides with the appearance of T-cell activation and proliferative markers in memory CD4^+^ T cells.** Untreated and 4 h or 24 h TCR-activated memory CD4^+^ T cells were subjected to immunofluorescence flow cytometry to monitor the expression of cyclinT1, pSer175 or pThr186 T-loop phosphorylated CDK9, the activated form of NF-κB (pSer529 p65 NF-κB), T-cell proliferative markers Ki67 and CyclinD3, and the T-cell activation surface markers CD25 and CD69.

**S3 Fig. Kinetic examination of P-TEFb, Ki67 and cyclinD3 expression in central and effector memory CD4^+^ T-cell subsets following TCR co-stimulation.** *(A)* and *(B)* Following purification of these memory subsets from healthy donor PBMCs using the EasySep™ Human Central and Effector Memory CD4^+^ T Cell Isolation Kit (Cat. # 17865), they were stimulated through the TCR with anti-CD3/anti-CD28 Dynabeads for varying times as shown. Thereafter, cells were subjected to immunofluorescence flow cytometry to monitor the expression of the P-TEFb (pSer175 CDK9 and cyclinT1), Ki67 and cyclin D3.

**S4 Fig. PKC agonists can sufficiently stimulate active P-TEFb expression in polarized primary Th17 cells.** *(A)* Procedure for generating polarized quiescent primary Th17 cells from naïve CD4^+^ T cells with or without a latent infection with HIV. *(B)* A representative immunofluorescence flow cytometry experiment showing that PKC agonists can sufficiently induce active P-TEFb expression in polarized primary Th17 cells while TNF-α and the HDAC inhibitor SAHA are poor stimulators. Active P-TEFb expression was assessed by examining the dual expression of cyclinT1 and pSer175 CDK9 using fluorophore-conjugated antibodies.

**S5 Fig. Reactivation of latent HIV in primary Th17 cells by TCR co-stimulation or PKC agonists is unlikely to be mediated by PKC.** *(A)* Rapid expression of active P-TEFb in memory CD4^+^ T cells in response to TCR co-stimulation. Memory T cells from three different healthy donors (color-coded) were activated or not for 2 h through the TCR with anti-CD3/anti-CD28 Dynabeads. Afterwards, the cells were subjected to flow cytometry analysis following intracellular immunofluorescence staining for P-TEFb (by co-staining for CycT1 and pSer175 CDK9) or the C-terminal domain Ser2 phosphorylated form of RNA polymerase II (pSer2 RNAP II CTD). *(B)* A selective inhibitor of CDK9 kinase, flavopiridol (FVP) effectively blocks TCR-mediated proviral reactivation in the QUECEL primary Th17 model of HIV latency. Latently infected Th17 cells were pretreated or not with 100 nM FVP for 30 min prior to TCR co-stimulation with anti-CD3/anti-CD28 Dynabeads for 24 h. Thereafter, cells were analyzed by flow cytometry following immunostaining using a fluorophore-conjugated antibody towards the HIV Nef accessory protein. Error bars denote S.E. of the mean from three separate experiments. Statistical significance (*p* values) in both *A* and *B* was calculated using a two-tailed Student’s *t* test. *(C)* TNF-α and SAHA are inadequately able to reactivate latent HIV in primary Th17 cells. Proviral HIV expression was examined by immunofluorescence flow cytometry following immunostaining using a fluorophore-conjugated antibody towards the HIV Nef accessory protein. *(D)* Ingenol, prostratin and PMA can sufficiently reactivate latent HIV in primary Th17 cells. *(E)* Representative flow cytometry data showing that a combination of two PKC inhibitors (Ro-31-8220 and Gö 6983) are ineffective at blocking proviral reactivation in response to treatment with 200 nM ingenol or 1 μM prostratin. *(F)* Viability assessment of cells treated or not with a combination of Ro-31-8220 and Gö 6983 at 100 nM each prior to TCR co-stimulation or challenge with PKC agonists. Cell viability was assessed by flow cytometry following staining with the eFluor450 viability dye. The viability data shown are expressed as a percentage relative to non-treated cells.

**S6 Fig. Immunoblotting and bulk RNA-seq analysis of P-TEFb, 7SK snRNP and SEC expression in primary CD4^+^ T cells.** *(A)* Immunoblotting analysis of resting and TCR-activated primary Th17 cells to examine the expression of P-TEFb subunits (CDK9 and CycT1), its 7SK snRNP inhibitory partner HEXIM1, SEC components (ELL2, AFF1 and ENL), and transcription factors (NF-κB, NFAT-c1 and AP1). Resting polarized Th17 cells prepared from healthy donor naïve CD4^+^ T cells were stimulated or not with anti-CD3/anti-CD28 Dynabeads for the times shown prior to preparation of low salt and chromatin nuclear fractions. Thereafter, both fractions were subjected to Western blotting using primary antibodies towards the proteins shown. *(B)*, *(C)* and *(D)* Examination of the mRNA expression of P-TEFb, SEC and 7SK snRNP subunits using a publicly available bulk RNA-seq dataset (SRA accession SRP026389) prepared using human memory CD4^+^ T cells that had been activated or not through TCR co-stimulation with anti-CD3/anti-CD28 coated beads for the times shown.

**S7 Fig. Single-cell transcriptome profiling of the expression of P-TEFb, SEC and 7SK snRNP subunits in memory CD4^+^ T cells.** *(A)* and *(B)* Healthy donor-derived memory CD4^+^ T cells were allowed to recover overnight in complete RPMI medium with no IL-2 addition prior to challenging them or not with anti-CD3/anti-CD28 Dynabeads for 24 or 72 h at a 1:1 bead-to-cell ratio in media containing 15 IU/ml IL-2. Thereafter, cells were prepared for Drop-seq by magnetically removing the T-cell receptor activator beads, removal of dead cells using the MACS dead cell removal kit and resuspension of cells in 1X PBS containing 0.1% BSA at 2 × 10^5^ cells/ml. Cells were clustered according to their unbiased transcriptome signatures (unsupervised clustering) and the dot plot function in Seurat was used to quantitate positive cell enrichment and relative expression levels of P-TEFb, SEC and 7SK snRNP transcripts in the dataset.

**S8 Fig. Pan-PKC inhibitors are ineffective at suppressing activation loop phosphorylation of CDK9 at Ser175 and Thr186 in memory CD4^+^ T cells.** Cells were treated or not for 30 min with a combination of Ro-31-8220 and Gö 6983 at 100 nM each prior to stimulation with anti-CD3/anti-CD28 Dynabeads or challenge with ingenol or prostratin. Afterwards, cells were analyzed by flow cytometry following immunostaining using fluorophore-conjugated antibodies towards pThr186 CDK9 and pSer175 CDK9.

**S9 Fig. Elevation of active P-TEFb expression in memory CD4^+^ T cells following TCR co-stimulation is not dependent on PKC activity.** *(A)* Scheme for the activation of the transcription factor NF-κB by PKC in primary T cells upon TCR co-stimulation. *(B)* Immunoblotting analysis of memory CD4^+^ T-cell extracts demonstrating that the pan-PKC inhibitor Ro-31-8220 can retard the degradation on IκB-α and the nuclear induction of NF-κB. Healthy donor memory CD4^+^ T cells were treated or not with 100 nM Ro-31-8220 for 30 min prior to stimulation with anti-CD3/anti-CD28 Dynabeads for the times shown. Thereafter, low salt and chromatin nuclear fractions were prepared and subjected to Western blotting using primary antibodies towards IκB-α and NF-κB subunits p50 and p65. *(C)* PKC inhibitors are unable to suppress the induction of active P-TEFb in memory CD4^+^ T cells. Cells were treated or not for 30 min with Ro-31-8220, Gö 6950, or Gö 6983 at 100 nM each prior to TCR co-stimulation. Thereafter, cells were analyzed by flow cytometry following immunostaining using fluorophore-conjugated antibodies towards cyclinT1 and pSer175 CDK9.

**S10 Fig. The MEK inhibitor U0126 effectively blocks the activation of ERK1/2 and the nuclear expression of c-Fos in T cells.** *(A)* Kinetic analysis of the activation of the MAP kinase isoforms ERK1 and ERK2, and the nuclear mobilization of c-Fos and NF-κB in response to PMA treatment with or without pretreatment with the MEK inhibitor U0126. Cytosolic and nuclear extracts were prepared from Jurkat T cells that were challenged or not with PMA for the indicated times in the presence or absence of U0126. These extracts were subjected to Western blotting to examine the expression of ERK1/2 and their phosphorylation by MEK, inducible expression of the transcription factor c-Fos, and the nuclear induction of c-Fos, NF-κB and NFAT proteins. *(B)* TCR activation of ERK1/2 phosphorylation by MEK in memory CD4^+^ T cells and subsequent ERK regulation of c-Fos are effectively blocked by U0126. Whole cell extracts were prepared from healthy donor memory CD4^+^ T cells stimulated or not through the TCR with anti-CD3/anti-CD28 Dynabeads for the times shown. These extracts were subjected to Western blotting to examine the expression of ERK1/2 and their phosphorylation by MEK, and inducible expression of the transcription factor c-Fos.

**S11 Fig. MEK inhibitor U0126 effectively blocks the formation of active P-TEFb in memory CD4^+^ T cells that is in response to stimulation with PKC agonists.** Healthy donor memory CD4^+^ T cells were treated or not for 30 min with U0126 prior to 24 h TCR co-stimulation or challenge with ingenol, prostratin or PMA. Afterwards, cells were analyzed by flow cytometry for active P-TEFb expression following immunostaining using fluorophore-conjugated antibodies towards cyclinT1 and pSer175 CDK9.

**S12 Fig. MEK inhibitor U0126 effectively blocks synthesis of cyclin T1 and formation of pThr186 CDK9 in memory CD4^+^ T cells in response to PKC agonists.** Healthy donor memory CD4^+^ T cells were treated or not for 30 min with U0126 prior to 24 h TCR co-stimulation or challenge with ingenol or PMA. Afterwards, cells were analyzed by flow cytometry for cyclin T1 and pThr186 CDK9 expression following immunostaining using fluorophore-conjugated antibodies towards cyclin T1 and pThr186 CDK9.

**S13 Fig. MEK inhibitor U0126 effectively blocks the formation of active P-TEFb in polarized primary Th17 cells that is in response to stimulation with PKC agonists.** Resting primary Th17 cells were treated or not for 30 min with U0126 prior to 24 h TCR co-stimulation or challenge with ingenol, prostratin or PMA. Afterwards, cells were analyzed by flow cytometry for active P-TEFb expression following immunostaining using fluorophore-conjugated antibodies towards cyclinT1 and pSer175 CDK9.

**S14 Fig. Representative experiment demonstrating the effect of MEK inhibitor U0126 on proviral reactivation in polarized primary Th17 cells in response to TCR-costimulation and PKC agonists.** Latently infected resting primary Th17 cells were treated or not for 30 min with U0126 prior to 24 h TCR co-stimulation or challenge with the PKC agonists ingenol, prostratin or PMA. Thereafter, cells were analyzed by flow cytometry following immunostaining using a fluorophore-conjugated antibody towards HIV Nef.

**S15 Fig. Effect of MEK and JNK inhibition on proviral expression in primary T cells.** *(A)* Proposed scheme for the regulation of proviral reactivation by ERK and JNK MAPK signaling in TCR-activated primary T cells. *(B)* U0126 performs better than the JNK inhibitor SP600125 at suppressing the reactivation of latent HIV by TCR co-stimulation in HIV-infected Th17 cells albeit with noticeably significant variability. Latently infected primary Th17 cells prepared using naïve CD4^+^ T cells from three different donors were pretreated or not with U0126 or SP600125 for 30 min prior to TCR co-stimulation with anti-CD3/anti-CD28 Dynabeads for 24 h. Cells were then analyzed by flow cytometry following immunostaining using a fluorophore-conjugated antibody towards HIV Nef. The *p* values shown were calculated using a two-tailed Student’s *t* test.

**S16 Fig. The MAPK ERK pathway is a primary mediator of proviral reactivation in TCR-activated primary T cells.** *(A)* Latently infected Th17 cells prepared using healthy donor naïve CD4^+^ T cells were treated or not for 30 min with either the MEK inhibitor U0126 or the JNK inhibitor SP600125 on their own or in combination prior to 24 h TCR co-stimulation or challenge with either ingenol or prostratin. Afterwards, cells were analyzed by flow cytometry following immunostaining using a fluorophore-conjugated antibody towards HIV Nef. The graph shows data from two separate experiments. (B) Viability assessment by propidium iodide staining of memory CD4^+^ T cells treated with U0126 or SP600125 on their own or in combination prior to 24 h TCR co-stimulation or challenge with either ingenol or prostratin. The viability data shown are expressed as a percentage relative to non-treated cells.

**S17 Fig. MAPK ERK is the primary MAP kinase that mediates phorbol ester-induced P-TEFb expression in memory CD4^+^ T cells.** Healthy donor memory CD4^+^ T cells were treated or not for 30 min with either the MEK inhibitor U0126 or the JNK inhibitor SP600125 on their own or in combination prior to 24 h TCR co-stimulation or challenge with ingenol. Afterwards, cells were analyzed by flow cytometry for active P-TEFb expression following immunostaining using fluorophore-conjugated antibodies towards cyclin T1 and pSer175 CDK9.

**S18 Fig. A Raf inhibitor AZ 628 effectively blocks the formation of active P-TEFb in memory CD4^+^ T cells that is in response to stimulation with PKC agonists**. Healthy donor memory CD4^+^ T cells were treated or not for 30 min with AZ 628 prior to 24 h TCR co-stimulation or challenge with ingenol or prostratin. Afterwards, cells were analyzed by flow cytometry for active P-TEFb expression following immunostaining using fluorophore-conjugated antibodies towards cyclinT1 and pSer175 CDK9.

**S19 Fig. Bulk RNA-seq analysis of the expression of RasGRP, Ras, Raf and Rap1 isoforms in primary QUECEL subsets and memory CD4^+^ T cells.** Quiescent CD4^+^ T cells that had been polarized into Th1, Th2, Treg and Th17 subsets using the QUECEL procedure were activated or not for 24 h with anti-CD3/anti-CD28 Dynabeads. Bulk RNA-seq datasets obtained using these cells were analyzed to examine the expression of RasGRP *(A)*, Ras *(C)*, Raf *(D)* and Rap1 *(E)* isoforms. A publicly available bulk RNA-seq dataset of primary human memory CD4^+^ T cells that had been activated or not through TCR co-stimulation with anti-CD3/anti-CD28 coated beads (SRA accession SRP026389) was also analyzed for these factors (Tm Expt 1 and Expt 2). In *B*, additional assessment of RasGRP isoform expression was performed over the entire time course of TCR activation (0, 2, 4, 6, and 24 h) for which the memory T-cell bulk RNA-seq dataset was generated. Transcripts per million (TPM) values were used to evaluate the relative abundance of transcripts under resting and TCR-activated conditions. Statistical significance (*p* values) was calculated using a two-tailed Student’s *t* test.

**S20 Fig. Transcriptome profiling of PKC, DGK and mTORC transcripts in primary QUECEL subsets and memory CD4^+^ T cells.** *(A)* scRNA-seq analysis of unstimulated and TCR-activated memory CD4^+^ T cells. Healthy donor-derived memory CD4+ T cells were stimulated or not with anti-CD3/anti-CD28 Dynabeads for 24 or 72 h prior to being subjected to Drop-seq. Cells were clustered according to their unbiased transcriptome signatures (unsupervised clustering) and the dot plot function in Seurat was used to quantitate positive cell enrichment and relative expression levels of PKC, DGK and mTORC transcripts in the dataset. *(B)* Bulk RNA-seq analysis of primary QUECEL subsets and memory CD4^+^ T cells. Quiescent CD4^+^ T cells that had been polarized into Th1, Th2, Treg and Th17 subsets using the QUECEL procedure were activated or not for 24 h with anti-CD3/anti-CD28 Dynabeads. Bulk RNA-seq datasets obtained using these cells were analyzed to examine the expression of PKC, DGK and mTORC transcripts. A publicly available bulk RNA-seq dataset of primary human memory CD4^+^ T cells that had been activated or not through TCR co-stimulation with anti-CD3/anti-CD28 coated beads (SRA accession SRP026389) was also included in the analysis (Tm Expt 1 and Expt 2).

**S21 Fig. Effect of inhibiting AKT, GSK3β and PTEN on TCR-induced P-TEFb expression in memory CD4^+^ T cells.** *(A)* and *(B)* Healthy donor memory CD4^+^ T cells were treated or not for 30 min with inhibitors towards AKT (Ipatasertib), PTEN (VO-OHpic), GSK3β (SB216763 or Tideglusib) or PI3K (LY294002) at the concentrations shown prior 24 h TCR co-stimulation. Afterwards, cells were analyzed by flow cytometry for active P-TEFb expression following immunostaining using fluorophore-conjugated antibodies towards cyclinT1 and pSer175 CDK9.

**S22 Fig. Representative flow cytometry experiment demonstrating the effect of treating polarized primary Th17 cells with U0126 or LY292004 on their own or in combination on TCR-induced P-TEFb expression.** Resting primary Th17 cells were treated or not for 30 min with inhibitors targeting either MEK (U0126) or PI3K (LY294002) or both at the concentrations shown prior to TCR co-stimulation for 24 h. Thereafter, cells were examined for active P-TEFb expression by flow cytometry following dual immunofluorescence staining for cyclinT1 and pSer175 CDK9.

**S23 Fig. Combined inhibition of MEK and mTORC kinases substantially suppresses TCR-mediated provirαl reactivation in primary Th17 cells.** *(A)* Latently infected Th17 cells prepared using naïve CD4^+^ T cells isolated from a healthy donor were pretreated or not for 30 min with the inhibitors shown prior to TCR co-stimulation for 24 h. Afterwards, cells were analyzed by flow cytometry following immunostaining using a fluorophore-conjugated antibody towards HIV Nef. *(B)* Assessment of the effect of combined inhibitor treatment on cell viability. Cell viability was assessed by flow cytometry following staining with the eFluor450 viability dye. The viability data shown are expressed as a percentage relative to non-treated cells.

